# Evaluating the boundaries of marine biogeographic regions of the Southwestern Atlantic using halacarid mites (Halacaridae), meiobenthic organisms with a low dispersal potential

**DOI:** 10.1101/794040

**Authors:** Almir R. Pepato, Teofânia H. D. Vidigal, Pavel B. Klimov

**Affiliations:** Departamento de Zoologia, Instituto de Ciências Biológicas, UFMG, Av. Antonio Carlos 6627, 31270-901, Belo Horizonte, Minas Gerais, Brazil; Tyumen State University, 10 Semakova Str., 625003 Tyumen, Russia; University of Michigan, Department of Ecology and Evolutionary Biology, 3600 Varsity Dr., Ann Arbor, Michigan 48108-2888 United States

**Keywords:** Marine Provinces, Marine Ecoregions, Phylogeography, Western South Atlantic, Halacaridae, *Agauopsis*, *Rhombognathus*, environmental niche modeling

## Abstract

**Aim:** We evaluated traditional biogeographic boundaries of coastal marine regions in SW Atlantic using DNA sequence data from common, rocky-shore inhabiting, marine mites of the genera *Agauopsis* and *Rhombognathus*, family Halacaridae.

**Methods:** We investigated geographic population genetic structure using CO1 gene sequences, estimated divergence times using a multigene dataset and absolute time-calibrated molecular clock analyses, and performed environmental niche modeling (ENM) of common marine mite species.

**Results:** *Agauopsis legionium* has a shallow history (2.01 Ma) with four geographically differentiated groups. Two of them corresponded to the traditional Amazonian and Northeastern ecoregions, but the boundary between the two other groups was inferred at the Abrolhos Plateau, not Cabo Frio. *Rhombognathus levigatoides* s. lat. was represented by two cryptic species that diverged 7.22 (multilocus data) or 10.01 Ma (CO1-only analyses), with their boundary, again at the Abrolhos Plateau. ENM showed that *A. legionium* has suitable habitats scattered along the coast, while the two *R. levigatoides* cryptic species differ considerably in their niches, especially in parameters related to upwelling. This indicates that genetic isolation associated with the Abrolhos Plateau occurred in both lineages, but for the *R. levigatoides* species complex, ecological niche specialization was also an important factor.

**Main conclusions:** Our study suggests that the major biogeographic boundary in the Southwestern Atlantic lies not at Cabo Frio but at the Abrolhos Plateau. There, two biogeographically relevant factors meet: (i) changes in current directions (which limit dispersal) and (ii) abrupt changes in environmental parameters associated with the South Atlantic Central Waters (SACW) upwelling (offering distinct ecological niches). We suggest that our result represents a general biogeographic pattern because a barrier at the Abrolhos Plateau was found previously for the fish genus *Macrodon* (phylogeographic data), prosobranch mollusks, ascidians, and reef fishes (community-level data).

## INTRODUCTION

Concordance of phylogeographic (population level) and biogeographic (above species level) patterns provide a useful framework for investigating and comparing patterns of evolution of faunas around the globe (Dawson, 2001). Such concordance is expected since faunal provinces are based upon endemism (Briggs, 1974), and the same factors (e.g. barriers to dispersal, divergent natural selection) that fuel speciation also lead to population genetic structuring, making both processes interchangeable in the short term (Nosil et al., 2009, Sukumaran & Knowles, 2017). As stated by the founders of phylogeography, “macroevolution is ineluctably an extrapolation of microevolution” (Avise et al., 1987).

In establishing those patterns, however, not all marine organisms are equally informative. There is a clear relationship between genetic structure, dispersal ability, fecundity and habitat. Organisms without a pelagic period usually demonstrate sharper genetic structuring (Kelly & Palumbi, 2010 and references therein). Low fecundity and a mid- or high-tidal habitat also promote genetic differentiation (Dawson, 2001). High fecundity and planktonic dispersive larvae are associated with slight if any genetic structure as observed in many studies performed along the Brazilian coast: mangrove crabs (Oliveira-Neto et al., 2007), land blue crabs (Oliveira-Neto et al., 2008), interstitial ribbon worms (Andrade et al., 2011) and littorinid snails (Andrade et al., 2003).

Counterintuitively, when associated with natural selection, strong genetic population structuring could be found even in species with long-term pelagic larvae, or even completely pelagic species. In these species, local adaptation (“local” often spans over hundreds of kilometers in the marine realm) could lead to speciation despite high immigration rates of poorly fit genotypes outside of the selective environment (Sanford & Kelly, 2011, Palumbi, 1992).

In the marine environment, breaking points in genetic structuring or community composition are commonly associated with: (1) hydrodynamics, such as currents, affecting recruitment and/or availability of planktonic dispersal stages (e.g. Francisco et al., 2018); (2) gradients in ecological factors that can impose a limit on geographic distributions or lead to local adaptation. However, they may be difficult to disentangle because changes in superficial circulation often create dispersal barriers where disparate water masses meet (Gaylord & Gaines, 2000, Pelc et al, 2009).

### Brazilian coastal bio- and phylo-geography

Spalding et al. (2007) classified the Brazilian coast into three biogeographic provinces: (1) North Brazil Shelf, ranging south to the Parnaíba River Delta; (2) Tropical Southwestern Atlantic extending to the upwelling area of Cabo Frio, Rio de Janeiro State; and (3) Warm Temperate Southwestern Atlantic, with its southern limit at Váldez Peninsula. These provinces were further subdivided into five ecoregions (areas of relatively homogeneous species composition, clearly distinct from adjacent systems) as shown on Fig. 1A. This division of coastal and shelf areas of the Brazilian coast greatly influenced subsequent studies on marine conservation and biogeography (e.g. Mantelatto et al., 2018, Huguenin et al., 2018, Bissoli & Bernardino, 2018). Importantly, this biogeographic classification adopted an older view that the main biogeographic break in the region is Cabo Frio, which is the southeast limit for coral reefs, and is in rough agreement with the 23°C isotherm data (Briggs, 1974, 1995, Palacios, 1982).

**Fig 1.**
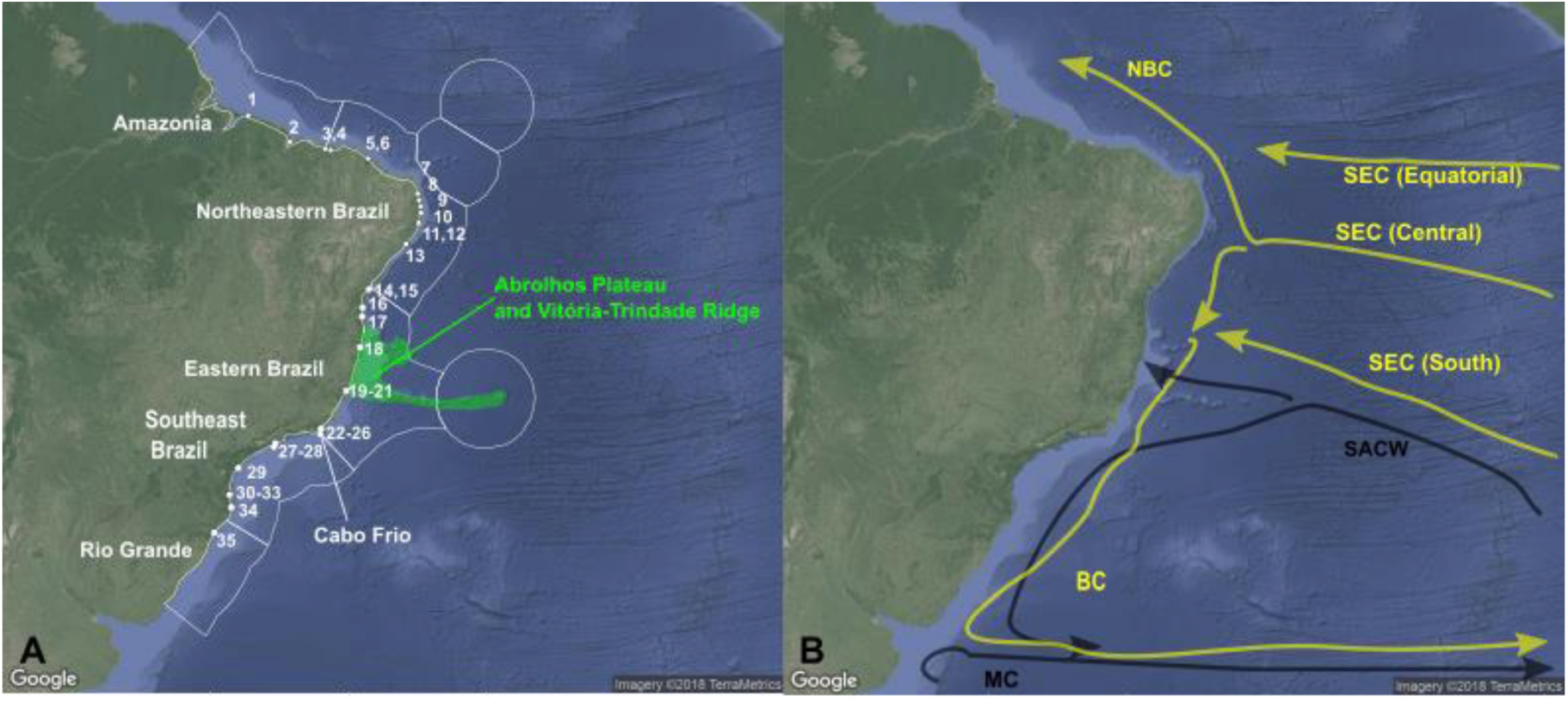
A - Map of Brazilian coast showing 36 sampling localities, traditional boundaries of ecoregions (Spalding et al., 2007). Abrolhos Plateau and Vitória-Trindade ridge are highlighted in green, Cabo Frio indicated. B - Southwestern Atlantic currents. Yellow arrows represent superficial warm, saline and oligotrophic currents, NBC: North Brazil Current; South Equatorial Current (SEC), with three branches: Southern (SSEC), Central (CSEC) and Equatorial (ESEC). Black arrows represent colder, less saline and more nutritive currents: the South Atlantic Central Water (SACW, 100-500 m depth), and Malvinas (Falklands) current (MC).

However, growing evidence suggests the need for modifications of this classification. Lotufo (2002), comparing ascidian communities, highlighted the role of the South Atlantic Central Waters (SACW) and the replacement of sandstone reefs by crystalline substrate starting from Bahia state (18° S) in establishing the southern limit for tropical fauna. Floeter et al. (2001) suggested that it lies in northern Espírito Santo (20° S), mentioning that Abrolhos reefs and the Vitória–Trindade Ridge form a topographical barrier to the Brazil Current inducing fundamental changes in physical, chemical and biological features over the SE shelf. Barroso et al. (2016) supported a similar conclusion with prosobranch gastropods, while Pinheiro et al. (2018) grouped the reef fish communities of Bahia with the Southeast. Similarly, coastal fishes of the genus *Macrodon* form two almost parapatric sister species (Santos et al., 2006), separated at the Abrolhos Plateau region (approx. latitudes 15-20°S): King weakfish *M. ancylodon* and Southern King Weakfish *M. atricauda* (Carvalho-Filho et al., 2010). Niche differentiation due to the SACW upwelling is the most plausible explanation for the presence of these two species, because, conceivably, these fishes can easily migrate over the Abrolhos Plateau region. In summary, overall data from nearly all available studies suggest that the southern limit for the tropical fauna should be northward of the traditional boundary at Cabo Frio.

### Main currents and coastal waters

Marine biogeographic boundaries usually occur where major oceanic currents interact. In the ocean, colliding currents may create flow-inducing boundaries affecting dispersal and distributional patterns near the shore (Gaylord and Gaines, 2000). Currents also determine the characteristics of Coastal Waters (CW), i.e. the outcome of interactions among oceanic water masses and continental drainage (Castro & Miranda, 1998), which directly influence intertidal animals (Fenberg et al., 2014).

The following regions have been defined by the coastal water characteristics along the Brazilian coast:

1. Amazon Shelf (4°N-2°S), where the huge discharge of freshwater from the Amazon River and other large rivers is an important parameter affecting CW, enriching the water with nutrients, increasing its turbidity and lowering its salinity (Castro & Miranda, 1998) ;
2. Northeastern (NES, 2° S-8 ° S) and Eastern (ES, 8° S-15 ° S) shelves, where the coast receives neither large inputs of freshwater nor the influence of the nutritive SACW (Castro et al., 2006, see below). This region is under the influence of the three warm, oligotrophic and saline branches of the westward flowing South Equatorial Current (SEC) that hit the Brazilian coast between 7°S and 17°S, directing most of its transport northward. The North Brazil Current (NBC) is formed south of 10°30’S, where the south and central SEC branches converge (Silveira et al., 1994, 2000, Fig 1B). From there, the NBC meets the northern branch of the SEC, turning northwestward. Just a minor portion of the water transported westward by the southern SEC branch turns south near 10°S, feeding the Brazil Current (BC). Between this latitude and the Abrolhos Plateau, local circulation and the southernmost branch of SEC that hits the littoral at approximately 17° S make the coastal water transport complicated, at some points with almost no transport or others with northward transport (Peterson & Stramma, 1991).
3. Abrolhos-Campos Region (ACR, 15°S-23°S) and the South Brazil Bight (SBB 23°S-28.5°S) are enriched by the diffusion of nutrients from SACW (Stramma & England, 1999). BC runs southwestward as far as 33°-38° S reaching the Subtropical Convergence (Castro et al. 2006, Peterson & Stramma, 1991, Stramma & England, 1999, Fig. 1B). The South Atlantic Central Water (SACW) has low salinities and temperatures, and high nutritive content. It results from subduction of surface waters at the Subtropical Convergence, flowing east and northeastward then being entrained into the South Equatorial Current and returning to the South American coast after receiving some water from the Indian Ocean (Reid, 1989; Stramma & England, 1999).

### Temporal context

As mentioned above, the AP region (Abrolhos Plateau and Vitória-Trindade ridge) can affect passive transport of marine organisms (e.g. Francisco et al., 2018; for theoretical models for colliding oceanic currents creating distributional patterns, see Gaylord & Gaines, 2000). At the same time, changes in the CW parameters would lead to local adaptation or to more or less permeable barriers. These changes also should take place at the AP region because the SACW influence limit occurs there. The presence of SACW is related to the presence of a fully operating cold Malvinas (Falkland) Current (MC), which in its turn is related to the opening of the Drake Passage. Despite superficial water circulation that may have started 36-28 Ma, the opening between South America and Antarctica became effective for deep waters not earlier than the middle Miocene (Crame, 1999). Martínez & del Río (2002), however, showed that mollusks from latitudes as south as 42 °S were inhabitants of warm waters by the late Miocene (Tortonian, up to 7.25 Ma), suggesting that the MC still had a very low activity in the area during that time. The minimal age for this system must correspond to the development of the West Antarctica ice sheet and intensification of oceanic circulation at the Miocene-Pliocene transition (4.8 Ma, Clarke & Crame, 1992).

### Halacarid mites as models for marine biogeography

This study aims at testing long held ideas about biogeographic division of the Brazilian coast using phylogeographic analyses of common marine meiobentic species as proxies. Those species were chosen because they have several natural history features maximizing divergence and maintaining genetic structure (Dawson, 2001): low fecundity, poor dispersal abilities and intertidal habitat.

Marine mites (family Halacaridae) differ from most common marine benthic organisms by the lack of a dispersing planktonic stage – immature mites look similar to adults and live in the same environment; the mites have low fertility and are unable to swim at any stage of their life cycle (Bartsch, 2004). They rely on passive means of dispersal, like rafting on algae or hitchhiking on large marine organisms covered by algae and debris (Bartsch, 2002). Halacarids have one larval and one to three nymphal stages (proto-, deuto-, tritonymph) before they molt to the adult stage. Although parthenogenesis is known in the family, they usually have separate sexes, with females slightly outnumbering males in a given population (Bartsch, 2006); this is the case in the two linages, *Rhombognathus* and *Agauopsis*, included in our study. Most species have a univoltine life cycle, with either short or prolonged periods of reproduction (Bartsch, 2006). In general, the fecundity is low, especially when compared to other marine invertebrates. Free-living species, like those studied here, produce less than 20 eggs per female lifetime (Bartsch, 2004, 2006).

Because halacarid mites are globally distributed, and are an ancient group diverging more the 300 Ma and radiating approximately 270 Ma from a terrestrial ancestor (Pepato et al., 2018), they can be used for inferring both ancient and recent biogeographic scenarios at local and global scales (Bartsch, 2004).

Halacarids are truly marine animals distinct from, for example, marine Oribatida, whose restriction to littoral zones suggests that they have not completely transcended the ecological barrier between marine and terrestrial environments (Procheş & Marshall, 2001). Halacarids inhabiting the upper intertidal fringe apparently evolved tolerance to desiccation and environmental fluctuations characteristic of such habitats (Pugh & King, 1985, Somerfield & Jeal, 1995). The family is exclusively benthic, with species living in interstitial spaces, on marine algae, or sessile invertebrates. Four halacarid genera feed on macroalgae. These were traditionally included in a single subfamily, Rhombognathinae, which included the genus *Rhombognathus* (Abe, 1998). In recent molecular studies, this subfamily was shown to be diphyletic (Pepato et al., 2018). In contrast to the algivorous *Rhombognathus*, species in another lineage included in our study, mites of the genus *Agauopsis*, were observed preying on copepods, ostracods and other halacarids, capturing and holding the prey between the femoral, genual and tibial crook and the spines of the first leg (Krantz 1970, MacQuitty, 1984).

### Hypotheses

Spalding et al.’s (2007) ecoregions are in rough agreement with some independently proposed divisions based on oceanographic data, e.g., breaks between Northeastern and Eastern ecoregions at 10° S and Northeastern and Amazonian ecoregions at 2° S. However, this view is at odds with the boundary between the Eastern and Southeastern regions. Spalding et al. (2007) placed this boundary at Cabo Frio (23°S), but the oceanographic data suggest that this boundary lies at the Abrolhos region (16 °S). Solving this controversy will be the main focus of our study. If the onset of current conditions took place at the Miocene-Pliocene transition, we expect that divergences of at least some sibling species could be relatively old, 4.8-7.5 Ma.

## Material and Methods

### Taxon sampling and sequencing

Sampling localities are reported in Fig. 1 and Appendix 1. We sequenced 609 bp of the mitochondrial gene Cytochrome Oxidase Subunit I (COI) for a total of 217 individual mites morphologically identified as *Rhombognathus levigatoides* Pepato & da Rocha, 2007 (n=128) and *Agauopsis legionium* Pepato & Tiago, 2005 (n=89). Collection data, GenBank accession numbers and details on amplification and sequencing are provided in Appendix 1. Close and distant outgroups were also included in our analyses (for detail, see below). *Rhombognathus levigatoides* was suggested to be a junior synonym of *Rhombognathus levigatus* Bartsch, 2000 (Abe & Fernandes, 2011) from the west coast of Australia (Bartsch 2000, 2003) based on the fact that the differences between the two taxa are slight (e.g. metric characters related to the first pair of dorsal setae and sclerotization). Irrespective of the status of *R. levigatoides* from Brazil, its populations lack morphological differences that would justify further splitting of this morphospecies.

To infer absolute time estimates of divergences of our target species (as detected by COI data) vs. closely related species, we included sequences of three genes [COI, small (18S) and large (28S) subunit ribosomal genes] from previous studies (Pepato & Klimov, 2015, Klimov et al., 2018, Dabert et al, 2016, Pepato et al., 2018) and a total of 23 terminals for which the sequences were generated as part of this study (Appendix 2, Table S1).

### Phylogeographic and population genetics analyses

Our COI alignment did not have indels or stop-codons, which are indicative of pseudogenes. Gene trees were inferred after choosing the best-fitting model and partition scheme in Partition Finder 1.0.1 (Lanfear *et al* 2012).

We computed haplotype features in DnaSP v. 5.10 and used Network 4.6.1.1 (2004–2012, fluxus-engineering.com) to construct median-joining networks (Bandelt et al, 1999). DnaSP 5.10 (Librado & Rozas, 2009) and MEGA7 (Kumar et al, 2016) were used to calculate population indices of haplotypic (Hd) and nucleotidic (π) diversity, and Arlequin 3.5 (Excoffier & Lischer, 2010) to calculate pairwise Φ-statistic (Φst) as a measure of genetic differentiation between different populations, with 10,000 permutations. For the later analysis, we employed the Tamura-Nei model because the best-fitting model obtained in jMODELTEST 2.1 was not available in Arlequin.

Alleles in Space (Miller, 2005) was used to correlate matrices of pairwise genetic distances (Φst) between all populations with geographic distances using a Mantel matrix correspondence test and produce scatterplots. Despite the limitations of this test in mtDNA analyses, usually lumping populations without pointing to Isolation by Distance in non-structured populations (Teske et al., 2018), we used the Mantel test to explore the data, interpreting the results according to Hutchinson & Templeton (1999).

SAMOVA 2 was employed to test our biogeographic hypotheses (Dupanloup et al, 2002). Our analyses were based on 1000 simulated annealing steps and the prior definition of the number of groups, K, ranging from 2 to 10. For each analysis with increasing K, we examined the proportion of genetic variance due to differences between groups, FCT, and searched the range of K for which FCT was the largest and statistically significant. We also calculated BIC (Bayesian Information Criterion) according to Meirmans (2012).

Additional AMOVA analyses were performed to compare our data-driven hypotheses with those constrained to reflect the Brazilian biogeographic provinces and ecoregions of Spalding et al. (2007).

Our COI phylogeny and haplotype networks (see below) suggested the presence of two distinct cryptic species in *Rhombognathus levigatoides*. To evaluate this result we conducted a barcoding gap discovery analysis in ABGD (Puillandre et al., 2012) and Generalized Mixed Yule Coalescent species delimitation analysis in bGMYC (Reid & Carstens, 2012).

For bGMYC analyses, all 99 haplotypes were included in a standard BEAST analyses employing the Yule process for modeling lineage divergence. Runs employing lognormal relaxed and strict clock models led to choosing the former due better fit according to AICM (ΔAICM = 6.18). We used a subsample of 500 *BEAST stationary trees; for each of these subsampled trees, we calculated species membership probabilities in the R package bGMYC (available at https://sites.google.com/site/noahmreid/software).

ABGD was run with the default values (web version), except by Nb bins = 64 and using Kimura (K80) Ts/Tv ratio = 4.304 as calculated in MEGA7. The sequences were directly submitted to the webserver instead of submitting a pre-calculated distance matrix.

### Absolute time estimation for mite divergences

A time calibration analysis was performed in BEAST (Bouckaert et al., 2014). We used a representative 169-taxon acariform taxonset, allowing time-calibration using paleontological data (12 calibration points, Appendix 2: Table 2). Outgroup sampling, calibration points, methodological details, and results of time-calibrated analyses using multiple loci are reported in Appendix 2.

Time calibrated COI gene trees were inferred in *BEAST, using the coalescent model and the species divergence priors informed by a separate 169-taxon, time calibration analyses based on fossils (see above, Appendix 2). For these secondary calibration points, we used lognormal priors (Morrison, 2008, Fig. 5) obtained by summarizing the absolute divergence times (see above) in TreeAnnotator 1.8.4, as mean and median, and setting the two values for the lognormal distributions in Beauti 2. This led to distributions with a variance slightly smaller than the original (that is multimodal, and hence difficult to represent as a single distribution), but still a good representation of uncertainties in time estimation.

**Fig. 2.**
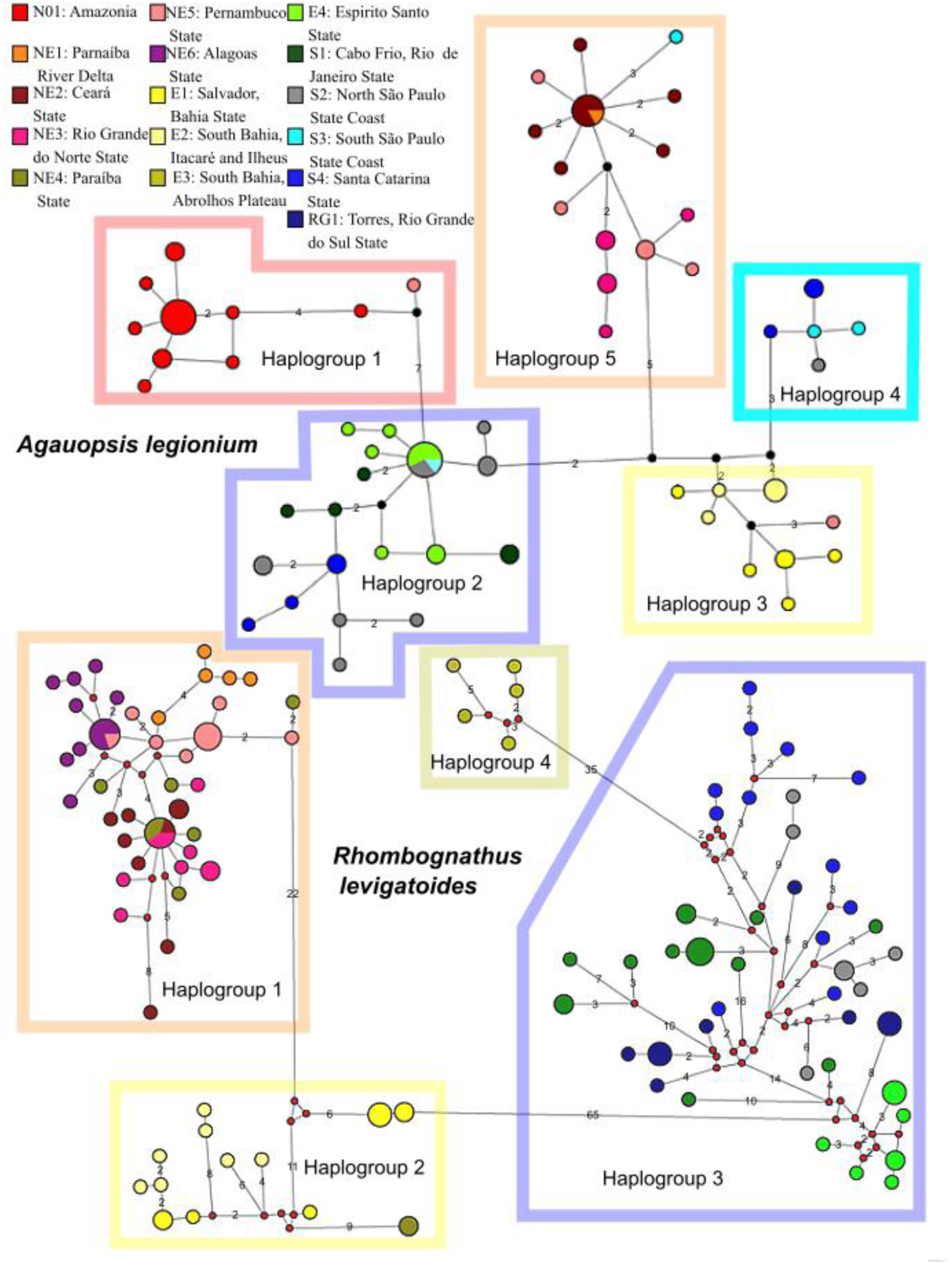
Minimum spanning network showing relationships among haplotypes of *Agauopsis legionium* and *Rhombognathus levigatoides*; different populations are color-coded. The number of mutational steps (>1) along the edges is indicated. Circles representing haplotypes are scaled to their frequencies. Haplogroups are referred to in the text.

**Fig. 3.**
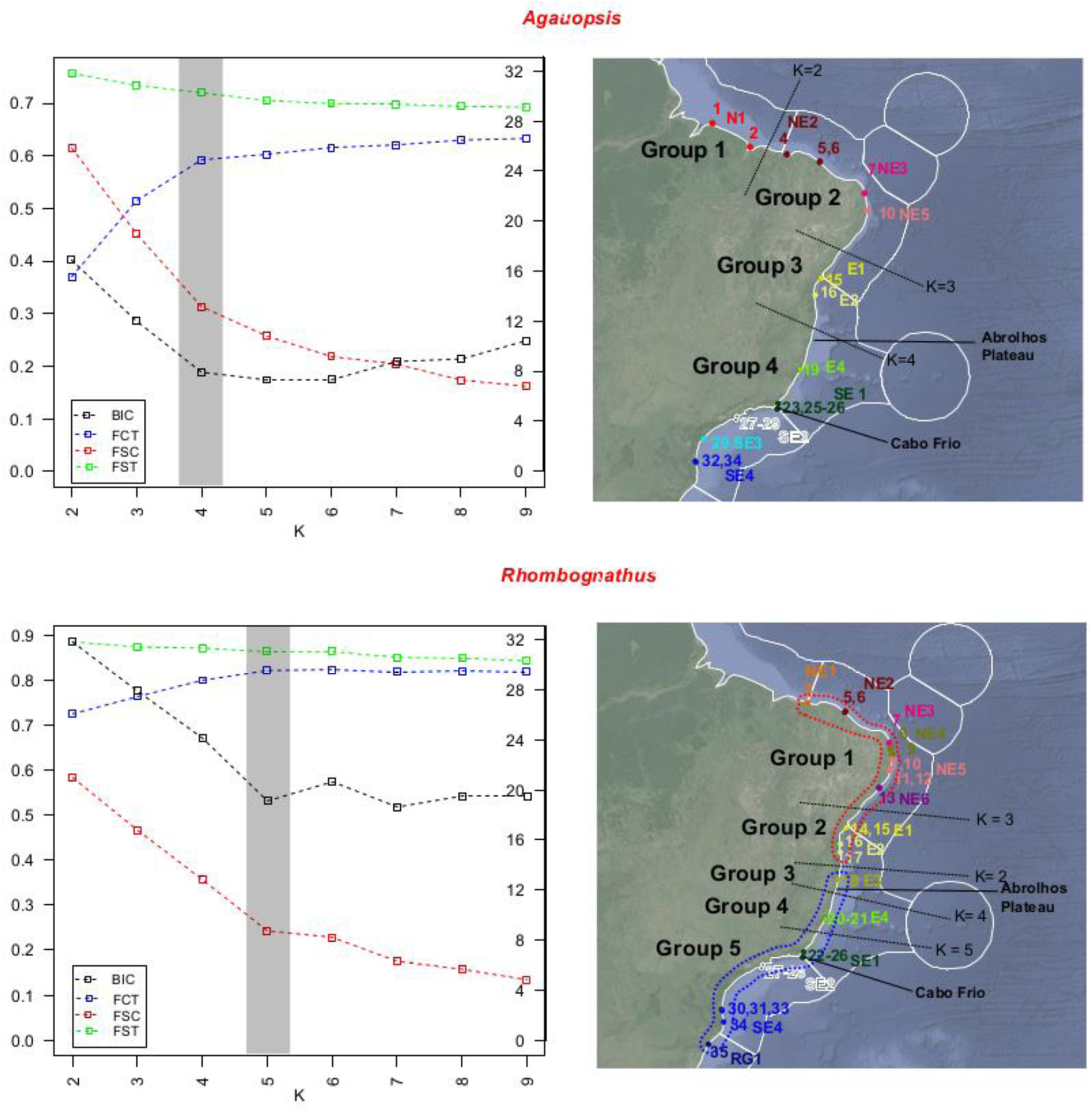
Left column: SAMOVA analyses with for K=2-9; shading indicates the optimal value of K; fixation indices FSC (among populations, within groups), FST (among populations), FCT (among groups) and Bayesian Information Criterion (BIC) are given. Left vertical axis scale for fixation indices, right vertical axis scale for BIC. Right column: geographic clusters recovered by SAMOVA; Sampling and traditionally recognized ecoregions (Spalding et al., 2007) are given; population color-coding follows that on Fig 2, it is indicated the Cabo Frio and the Abrolhos Plateau.

**Fig. 4.**
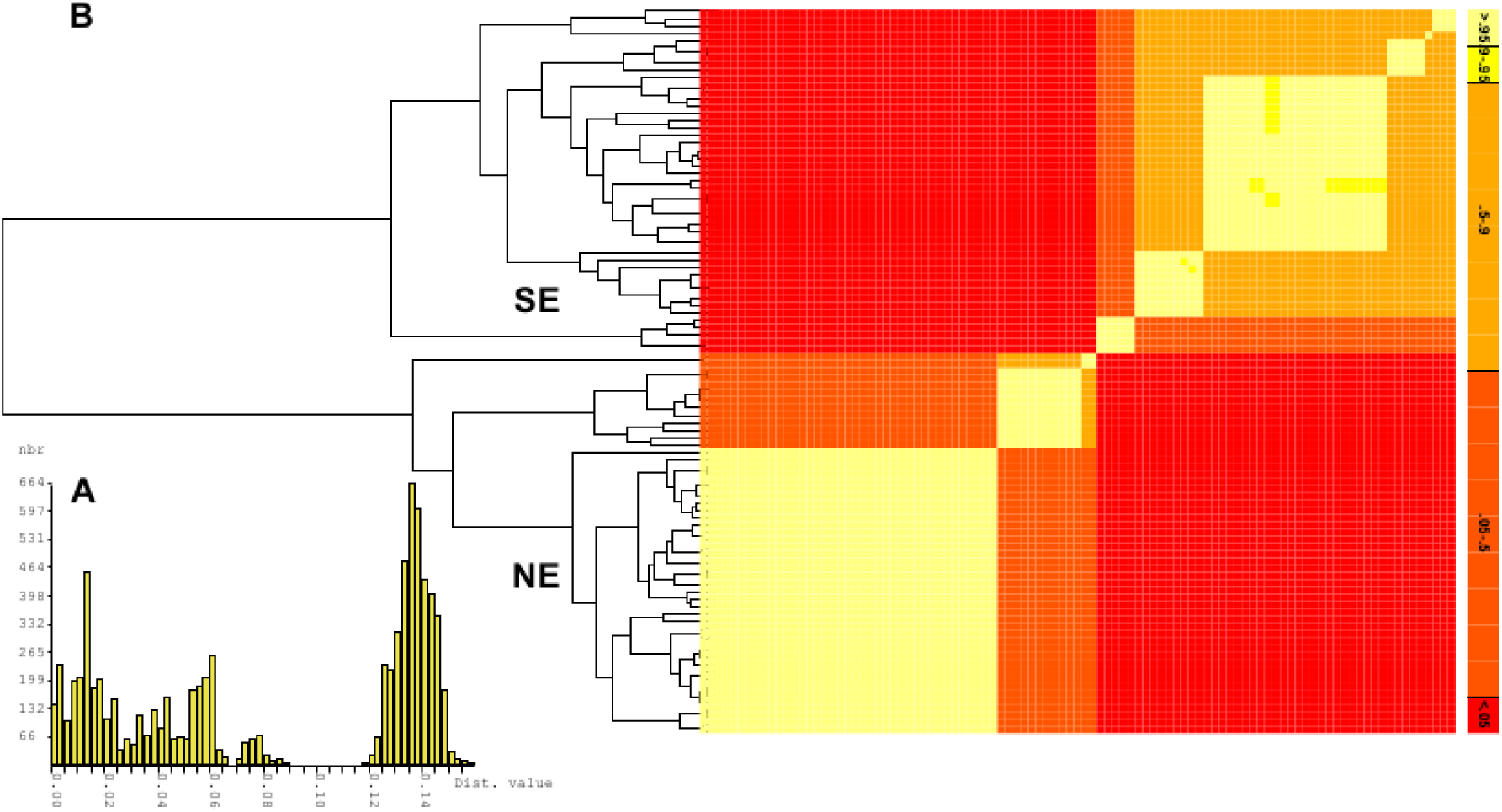
Species delimitation analysis of *Rhombognathus levigatoides* complex (ABGD) based on CO1. A. Gap between intra- and interspecific pairwise genetic distances; Dist. = K2P pairwise genetic distances, nbr= number of ranks. B. Maximum sum of clade credibility along with heatmap showing posterior probabilities for individuals to belong to the same species (i.e., evolve under the coalescent process). Note that the posterior probability for the NE and SE clades to be the same species is less than 0.05.

**Fig. 5.**
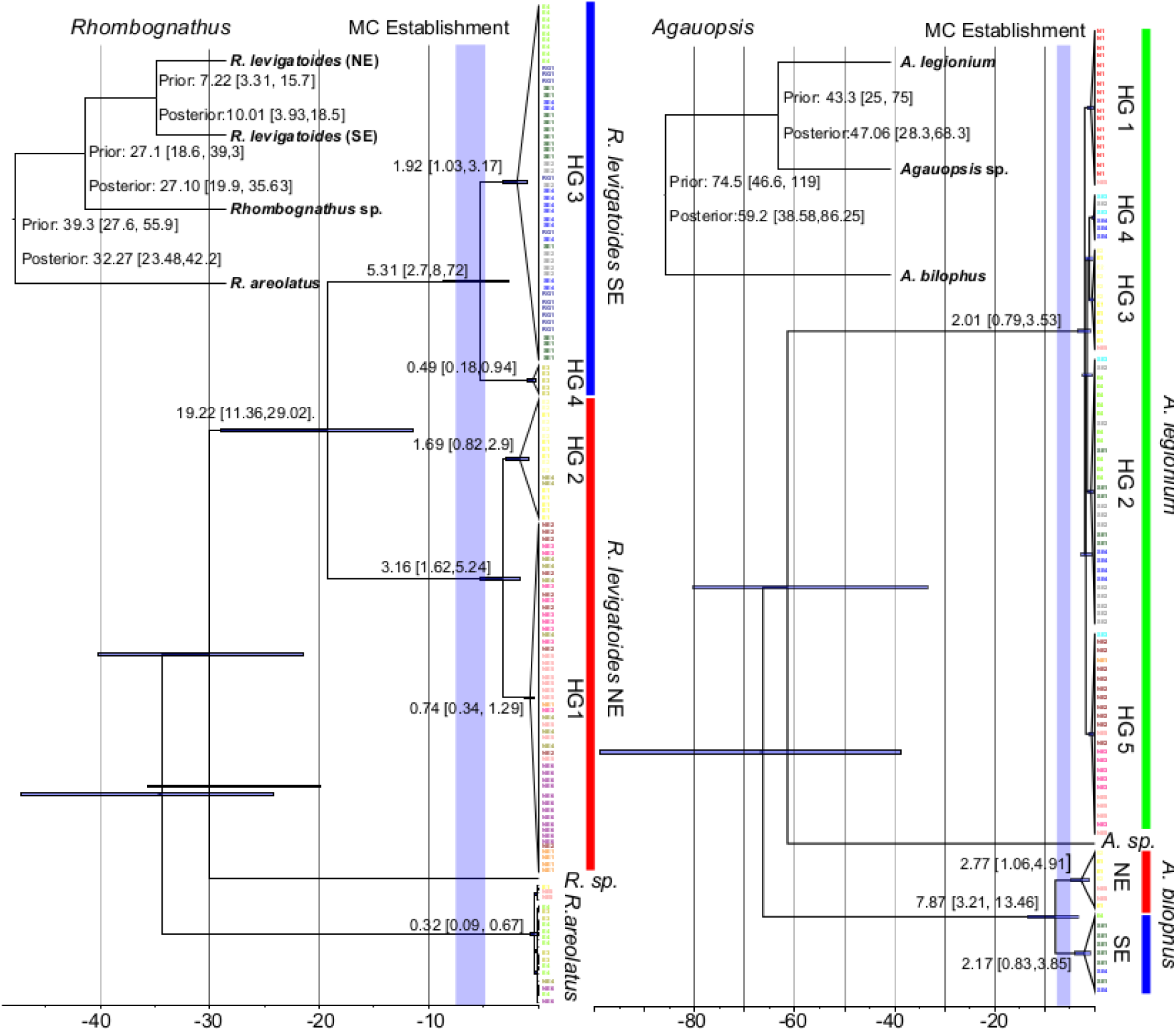
Time trees inferred from COI sequences under the multispecies coalescent model in *BEAST for *Rhombognathus levigatoides* and *Agauopsis legionium*. Numbers above branches are coalescence time estimates. Node bars represent 95% HPD intervals for time estimates. Vertical gray column indicates approximate period when the Malvinas (Falkland) current (MC) was established 7.25-4.8 Ma. Inserts are the species trees, indicating the prior (from the multilous analyses) and the posterior on divergence ages. HG, Haplogroups as referred in Fig. 2. Color code as in Figs. 2-3.

Strict and lognormal relaxed clocks were compared using AICM. The strict clock had a better fit compared to the relaxed lognormal in both *Agauopsis* (ΔAICM = 12.89) and *Rhombognathus* (ΔAICM = 62.45). Secondary calibration value estimates (median and mean) are given in Fig. 5.

### Species distribution modeling

Georeferenced data for the two species of the *Rhombognathus levigatoides* complex and *Agauopsis legionium* were employed as input for species distribution modeling in Maxent 3.4.1 (Phillips et al. 2006) and niche comparisons and modeling in the following R packages: ENMeval (Muscarella et al. 2014), ENMTools (Warren et al, 2010), and Ecospat (Di Cola et al., 2017). Maxent relies on the occurrence data to infer the niche parameters that shape species distributions (Phillips et al. 2006). Our R scripts are provided in Appendix 4.

We obtained 77 environmental layers from the Bio-ORACLE website (www.oracle.ugent.be/) (Tyberghein et al. 2012, Assis et al., 2017), MARSPEC (http://www.marspec.org/) (Sbrocco & Barber, 2013) and ECOCLIMATE (http://www.ecoclimate.org/) (Lima-Ribeiro et al., 2015). Because Bio-ORACLE and MARSPEC have 5 arcminute resolution and ECOCLIMATE 30 arcminute resolution, we employed the program DIVA-GIS to pre-process the layers for downstream analyses, to make the resolutions compatible by dividing ECOCLIMATE grid cells (Hijmans et al., 2001). The layers comprise oceanic (e.g. salinity, bathymetry) and atmospheric (e.g. precipitation) parameters since our target organisms are intertidal and may be influenced by non-marine environmental factors at low tide.

At the first step, the bathymetric layer from MARSPEC was employed to mask other layers in ArcMap 10, restricting their area to a strip along the coast shallower than 300 meters. Correlated environmental variables (i.e., absolute values of Pearson correlation coefficient larger than 0.75) were removed in ENMTools, resulting in 24 variables (Appendix 4, excluded highly correlated variables also shown).

We employed the 24 variables in pre-modeling niche comparison – i.e. a comparison made without inferring a model that expresses a probability distribution – between the two *Rhombognathus* sibling species, using the R package Ecospat (Di Cola et al., 2017). In this case, values were extracted from the environmental layers based on the occurrence points and a background withdrawn from a radius around them (20 Km). Principal Components Analysis (PCA) calibrated on the environmental background (PCA-env, Broennimann et al., 2012) was employed to measure and display graphically the niche overlap.

Schoener’s D metrics was used for niche comparisons (Schoener, 1970); this varies from 0 (no overlap) to 1 (complete overlap). Warren et al. (2008)’s niche equivalency tests were run taking D metrics as reference. It takes the presence points of species under consideration and randomly reassigns them to each species, then it checks if the species niches are drawn from the same underlying environmental parameter distribution. If the observed value of D falls within the density of 95% of the simulated values, the null hypothesis of niche equivalency cannot be rejected. To perform these tests, we ran 100 replicates in Ecospat (Di Cola et al., 2017).

Because of the relatively sparse sampling, we were careful to balance the model goodness-of-fit and complexity by tuning Maxent settings of feature classes and regularization multiplier. Two different feature classes (FC, linear and quadratic) and six values for regularization multiplier (RM, 0.5, 1.0, 1.5, 2.0, 2.5, and 3.0) were tested using the AICc in the R package ENMeval (Muscarella et al. 2014). By default, MAXENT automatically determines which FCs to use based on the number of occurrence localities (up to two in our case) and RM is set to 1. Data were partitioned using the n-1 jackknife method, in which each occurrence is used for testing, while all others are used for training in that iteration (Muscarella et al. 2014). The program was run iteratively: first, all layers selected by excluding highly correlated variables were used, but those variables with no contribution to the model along the range of settings examined were pruned. Models with the lowest AICc value and those differing by less than 2 (Δ AICc < 2) were considered as equivalent. If more than one set of parameters lie in this interval of AICc values, the choice was based on the maximum value of AUC test, i.e. on our ability to predict the distribution of independent occurrences in the training area (Warren & Seifert, 2011). These two criteria are based on different theoretical approaches; the former is rooted in Information Theory and model fit to the data, and the latter is based on the performance of independent test data. We consider these criteria complementary to each other.

Moreover, we ran analyses taking into account the effect of sampling bias by feeding into Maxent a bias file in which the cell values reflect sampling effort and give a weight to random background data used for modeling (Elith et al., 2010). The bias file was created by deriving a two-dimensional kernel density estimate, based on the coordinates of all sampling localities, using the R package MASS. An R script employed to construct the bias file is also provided in Appendix 4.

Because of the relatively sparse sampling, we ran MaxEnt in 10 replicates with cross-validation. Regularization and features set according ENMeval results, and presenting results as cloglog, which estimates the probability of the species presence across the grid (Merow et al., 2013).

## Results

### *Agauopsis legionium* phylogeography

The longest edge in the network (Fig. 2) had seven mutational steps, separating a group that comprises haplotypes from the Amazonian region (from Pará to Maranhão, 0°35’to 2°29’ S) plus a single haplotype from the Northeast (Pernambuco State, approx., 8°S) (Fig. 2: Haplogroup 1) from a group including haplotypes from Santa Catarina to Espirito Santo States (Fig. 2: Haplogroup 2, 19°56 to 26°48’S). The other three haplogroups comprise haplotypes from Bahia State (Haplogroup 3, 13° 1’ to 14° 17’ S) and a single haplotype from Pernambuco State; a small group with haplotypes from Santa Catarina and São Paulo States (Haplogroup 4, 23° 49’ to 26° 48’S); and a haplogroup composed mostly by Northeast samples (Haplougroup 5, 2° 55’ to 7° 38’ S), plus a single sample from south of São Paulo.

One-way AMOVA based on Tajima and Nei distances indicated strong differentiation among populations (Фst = 0.68, P < 0.001). Pairwise comparisons among populations failed to find significant values of Фst among the population pairs SE1/SE2, SE1/SE4, SE2/SE4, SE3/SE4, but the remaining pairwise combinations were significant. SAMOVA results showed an asymptotic behavior, with the last considerable raise of FCT and reduction of FSC values at K=4; the same result was obtained using BIC values (Fig. 3A). This suggests the following grouping scheme: Group1: Amazonian region, comprising samples from Pará and Maranhão States (Population N01); Group 2: Northeast from Piauí to Pernambuco (populations NE2, NE3, NE5); Group 3: Bahia State (populations E01, E02) and Group 4: Southeast from Espirito Santo to Santa Catarina States (populations E04, SE1, SE2, SE3, SE4). The order in which the groups were separated with the stepwise increase of K was: K=2, Group 1 vs Groups 2-4; K=3, Group 2 vs groups 1, 3-4.

In the preferred K=4 scheme, 59.23% of the variation was observed among groups, 12.77% among populations within groups, and 28.00% within populations. This contrasts with a scheme constrained to the traditional subdivision of marine provinces. AMOVA analyses including population E04 along E01 and E02 accounted for 48.34% among group variation, 22.23% among populations within groups, and 29.44% within populations. The scheme with the Abrolhos Plateau being the border between the East and Southeast regions was clearly favored over the traditional scheme placing the border at Cabo Frio (BICs 7.89 vs 13.44, respectively).

A Mantel test for the entire set of sequences indicated significant correlation between genetic and geographic distances (r=0.656, P<0.01; Fig. S3). The observed pattern suggests that the observed correlation is not related to isolation by distance.

### Rhombognathus levigatoides *phylogeography and species delimitation*

Our haplotype networks analysis revealed four haplogroups, all separated by a large number of mutational steps. The longest edge separated Haplogroup 2 (Populations E01 and E02, 13°0’-14°55’ S), from Bahia State, and two sequences from Paraiba State (6°41’ S) from the large Haplogroup 3 distributed from Espirito Santo to Rio Grande do Sul (19° 56’ S to 29° 21’S). Haplogroup 1 comprises most sequences from the Northeast, and is connected to Haplogroup 2 by 22 mutational steps. Finally, the small Haplogroup 4 was found in Cumuruxatiba, Bahia State (17° 4’ S, 39° 10’ W) and is connected to Haplogroup 3 by 35 mutational steps.

One-way AMOVA based on Tajima and Nei distances indicated strong differentiation among populations (Фst = 0.83, P < 0.001). Pairwise comparison among populations found significant values of Фst for all population pairs, except for NE3/SE2, NE1/NE4, and NE2/NE4. SAMOVA showed an asymptotic behavior, with the last considerable raise in FCT and reduction of FSC and BIC values at K=5 (Fig. 3). The following groups were inferred: Group 1, Northeast from Piaui to Alagoas State (populations NE1-NE5); Group 2, Bahia State (populations E01-E02); Group 3, South Bahia (Population E03); Group 4, Espirito Santo State (population E04); Group 5 from the Rio de Janeiro to Rio Grande do Sul States (populations SE1 to RG1). The order in which the groups were separated with the stepwise increase of K was: K=2, Groups 1+2 vs Groups 3-5; K=3, Group 2 vs groups 1,3-5; and K=4, Group 3 vs groups 1,2, 4-5.

The preferred biogeographic scheme explained 82.20% of the among-group variation, 4.32% among populations within groups, and 13.48% within populations. A scheme constrained to reflect the traditional subdivision of East and Southeast regions at Cabo Frio explained 72.55% of among groups variation, 16.02% among populations within groups, and 11.43% within populations. Again, the preferred scheme was substantially better than the constrained one (BIC = 24.15 vs 37.14, respectively).

The Mantel test resulted in considerable correlation between genetic and geographic distances (r = 0.772, P<0.01), with a pattern that does not imply isolation by distance. Because this result may be attributed to the presence of two distinct cryptic species (see below), a Mantel test was repeated for each of them (Fig. S3). For the clade distributed from Piauí to Bahia (populations NE1-NE5, and E01-E02), the correlation was r = 0.531 (P<0.01), while for the group comprising populations E03, E04, SE1 to RG1 the correlation was r = 0.426 (P<0.01). Even after considering the two species (see below), there was no evidence suggesting that the observed correlation is related to isolation by distance.

*Rhombognathus* species delimitation analyses with bGMYC and ABGD agree in inferring the two allopatric clades (see above) as separate species: There was a clear gap at p=0.90-0.12 attributed to the gap between interspecific and intraspecific distances, and bGMYC analyses resulted in two distinct species, with the posterior probability for the alternative, single-species delimitation being p < 0.05 (Fig.4). Hereafter, these species are referred to as NE and SE species of the *Rhombognathus levigatoides* species complex.

### *Coalescent times among* Rhombognathus *and* Agauopsis *species haplotypes*

*Agauopsis legionium* has a relatively recent history, originating 2.01, 0.79-3.53 Ma (median, 95% HPD), with Haplogroup 1 being basal and the remaining haplogroups recovered as follows: (Haplogroup 5, (Haplogroup 2, (Haplogorup 3, Haplogroup 4))) (Fig. 5B). It is interesting that the outgroup species, *Agauopsis bilophus*, shows a strong geographic structure, between northeast and southeast populations similarly to those of the *Rhombognathus levigatoides* complex (see below). Coalescence between the two *Agauopsis bilophus* clades occurred 7.87, 3.21-13.46 Ma, and the ingroup coalescence occurred at 2.77 1.06-4.91 Ma for the Northeast clade and 2.17 0.83-3.85 for the Southeastern clade.

The *Rhombognathus levigatoides* species complex has a deeper history than that of *A. legionium* (Appendix 2, Fig. 5). The divergence between the SE and NE species occurred 7.22, 1.69-16.28 (multilocus analysis) or 10.01, 3.93-18.5 Ma (COI analysis). The latter analysis inferred coalescence between the two groups at 19.22, 11.36-29.02 Ma; coalescence of the NE species occurred at 3.16 1.62-5.24 and for the SE species 5.31, 2.7-8.72 Ma. The outgroup, *R. areolatus*, had a very shallow gene phylogeny, displaying no apparent structuring.

### Pre-modeling comparison among niches

Schoener’s D metrics revealed a considerable overlap between the *A. legionium* niche and those of *R. levigatoides* (NE species D = 0.427 and SE species D = 0.326), contrasting with a very small overlap between the *R. levigatoides* niches, D = 0.072 (Fig. 6 A-C). It is also evident that *A. legionium* has a niche breadth much wider than both *Rhombognathus* SE and NE species. The niche equivalency test falls within the 95% confidence interval of the null distribution, except for the comparison between the sibling species of *Rhombognathus* (Fig 6 D-E).

**Fig. 6.**
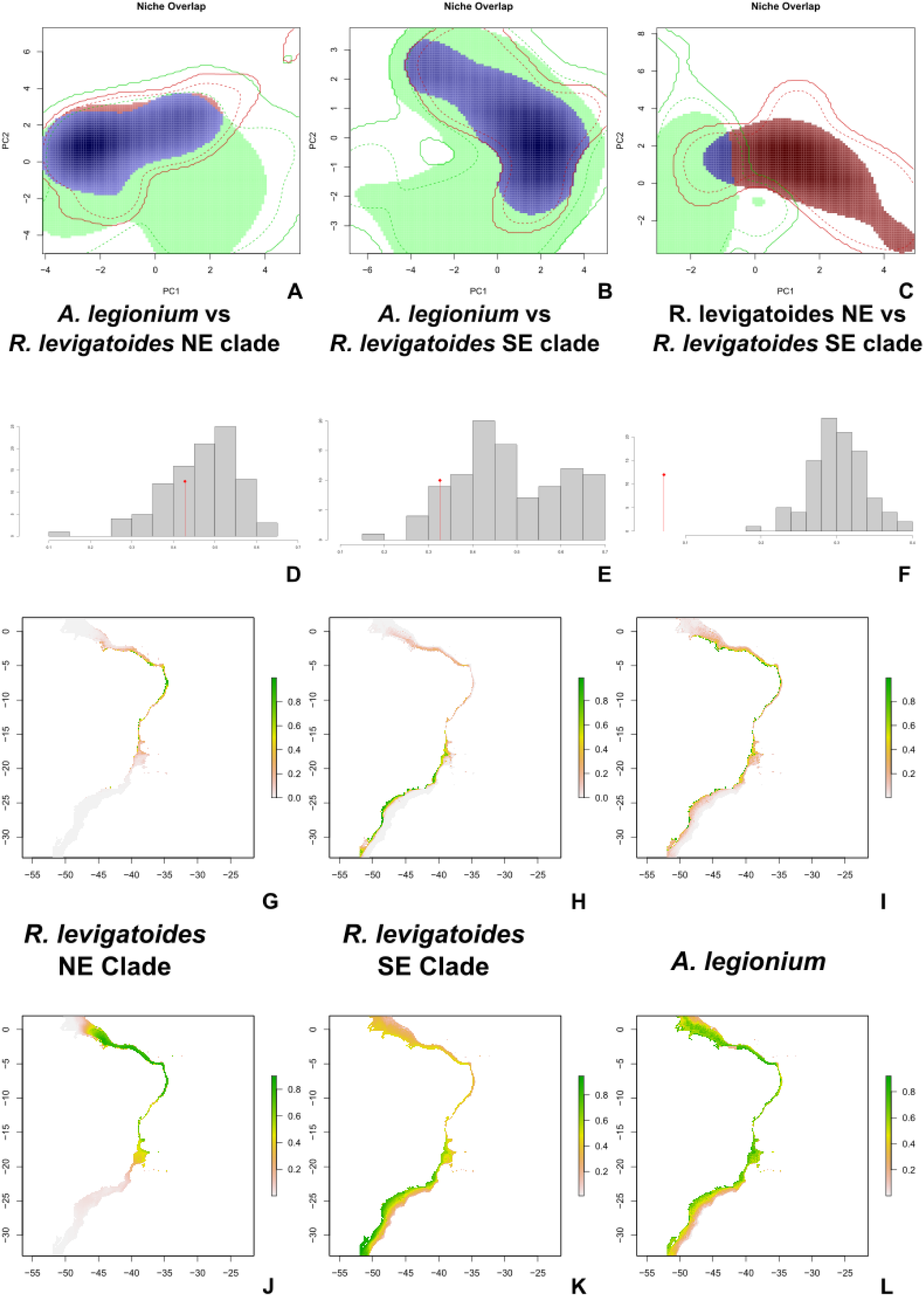
A-C: Representation of pairs of species niches along the two principal components of the PCA-env in: A- *Agauopsis legionium* and *Rhombognathus levigatoides* northeast species; B- *Agauopsis legionium* and *Rhombognathus levigatoides* southeast species; C- Both *Rhombognathus levigatoides* species. D-F Test of niche equivalence among the tree lineages, in the same order as PCA-env. The similarity score from the true data (red vertical line with a diamond) is plotted against the distribution of similarity scores built from occurrences drawn randomly in 100 replicates. The vertical axis is the frequency for each score interval. Distribution models inferred by MaxEnt ignoring (G-I) and accounting for (J-L) potential sampling bias. Latitude and Longitude are the vertical and horizontal axis. Color scale refers to suitability.

Correlation of variables with PCs is summarized in Appendix 4, Table 2. When analyzing the two species of *Rhombognathus*, the first two principal components explained 23.92% and 17.53% of the total variability, respectively. The following variables were highly correlated to the first PC: Sea Surface Temperature mean (−0.91), Chlorophyll A mean (−0.86), and Sea Surface Salinity of the freshest month (−0.85). The second PC is correlated with Bathymetry (−0.71), Annual precipitation (−0.69), and Minimal light at the bottom (−0.67).

### Species distribution modeling

Models accounting for sampling bias consistently had smaller AUC values and predicted distribution ranges broader as compared to models that do not incorporate sampling bias. These two main sets of models also pointed to different variables as main predictors.

The Northeast species of the *R. levigatoides* complex had its predicted distribution mainly over the Northeastern (NES, 2°S-8°S) and Eastern Shelf (ES, 8°S-15°S), irrespective if sampling bias was accounted for or not (Fig. 6 G and J, respectively). For the first model, ENMEval selected the use of linear features only and RM was 0.5. Model average AUC was 0.946. The variables with effective contribution to the model were Minimal light at the bottom (50.6%), Sea Surface Salinity of the freshest month (18.4 %), Mean diurnal temperature range (7.7%), Maximal current velocity (7.4%), Mean Chlorophyll A (4.6 %), Calcite concentration (4.4%); Mean sea surface temperature (3.8%), Maximal photosynthetically active radiation (2.1 %), and Maximal diffuse attenuation (1.1 %). Jackknife resampling identified the variable with the highest gain when in isolation to be Minimal light at the bottom (i.e., this variable provides the most useful information by itself), and the environmental variable that decreases the gain the most when omitted was Sea Surface Salinity of the freshest month (i.e., this variable provides the most information that is not present in other variables). When the sampling bias was taken into account, both AICc and AUC tests suggested linear and quadratic features, and RM was 1. Model average AUC was 0.862. The variables that contributed the most were: Mean sea surface temperature (65%), Calcite concentration (13.4 %), Sea Surface Salinity of the freshest month (7.7 %), Minimal light at the bottom (7.4 %), Mean concentration of Chlorophyll A (4.4 %), and bathymetry (2.1%). The jackknife test of Maxent of variable importance identified the variable Mean sea surface temperature as a variable providing both the highest gain (when used in isolation) and decreasing the gain the most (when omitted).

The Southeast species was predicted to be distributed at the Abrolhos-Campos Region (ACR, 15°S-23°S), South Brazil Bight (SBB 23°S-28.5°S) to Southern Shelf (SS, 28.5°S-34°S) (Fig. 6 H and K). When potential sampling bias was not accounted for, ENMEval suggested using linear and quadratic features, and RM was 1.5. Resulting model average AUC was 0.960. The variables that contributed the most were Bathymetry (52.3 %), Mean sea surface temperature (20.4%), Mean Calcite concentration (9.7%), Maximal Current velocity (7.4 %), Sea surface salinity of the freshest month (4.7%), Mean diurnal temperature range (3.6%), and Minimal light at the bottom (2.1%). The highest ranked variable was Bathymetry, showing the highest gain (when used in isolation) and also decreased the gain the most (when omitted). When the sampling bias was accounted for, AICc and AUC tests suggested using linear features only; RM was 2.0, and AUC was 0.864. The variables that contributed the most to the model were Sea Surface Temperature mean (76.6 %), Bathymetry (14.9%), and Maximal Current velocity (8.6%). Mean sea surface temperature was the highest ranked variable by the jackknife test.

Both *Agauopsis legionium* distribution models predicted multiple suitable localities scattered along the coast (Fig. 6 I and L). Without considering sampling bias, ENMeval suggested using linear features only, and RM was 0.5. The model AUC was 0.846, with the major contributions from the following three variables: Maximal light at the bottom (61.1%), Bathymetry (16.1), Sea Surface Salinity of the freshest month (7.7%), Minimal light at the bottom (5.6%), Minimal Nitrate concentration (4.2%), North/South Aspect (2.0%), Sea Surface Temperature mean (1.5%). The environmental variable with highest gain (when used in isolation) was Maximal light at the bottom, whereas Bathymetry decreased the gain the most (when omitted from the model). When sampling bias was accounted for, linear and quadratic features were suggested; RM was 1.5. The model AUC was 0.737, and the largest contribution was from the following variables: Maximal light at the bottom (37.4%), Precipitation of coldest quarter (22.3%), Bathymetry (20.5%), Minimal light at the bottom (7.6%), East/West Aspect (7.4%), and Maximal Current velocity (4.7%). Precipitation of coldest quarter was the highest ranked variable by the jackknife test.

## Discussion

In the marine environment, two key aspects may influence the presence of genetic structuring or lack thereof: (1) hydrodynamics, such as currents, affecting recruitment and/or availability of passively transported propagules, and (2) gradients in ecological factors that can impose a limit on geographic distributions (Andrade et al., 2011). The halacarid mite populations studied here do show a remarkable genetic structuring along the Brazilian coast. In contrast, a substantial number other coastal organisms inhabiting the same area show slight if any genetic structure. For example, mangrove crabs, *Ucides cordatus* (Oliveira-Neto et al., 2007), land blue crabs (Oliveira-Neto et al., 2008), interstitial ribbon worms, *Ototyphlonemertes erneba* and *O. evelinae* (Andrade et al., 2011) and littorinid snails, *Nodilittorina lineolata* and *Littoraria flava* (Andrade et al., 2003). Marine mites lack any long-distance dispersal stage and arguably do not extensively engage in passive dispersal by rafting on drifting algae (Bartsch, 2004). Therefore, the presence of considerable geographic genetic structuring in both species of marine mites is not surprising.

Populations of *A. legionium* and the two allopatric species of the *R. levigatoides* complex share a common break point at the Abrolhos Plateau (AP). In addition, ecological niche modeling demonstrated only a small niche overlap for the NE and SE species of the *R. levigatoides* complex. We explain these patterns by the fact that at the AP region, two biogeographically important factors meet: (i) a physical barrier (change of current direction) that affects connectivity between populations, and (ii) sharp changes in ecological parameters due to the SACW upwelling system, which extends south from AP along the Brazilian coast (for detail see the Introduction section). The distribution pattern observed in *A. legionium* would be related only to the first factor (physical barrier) because our environmental niche modeling analysis indicates that this species is not sensitive to the environmental gradients associated with SACW. However, the distributions of the two NE and SE species of the *R. levigatoides* complex are affected by the presence or absence of SACW upwelling southward to AP (see below). This result is congruent to the observations made for rock-shore organisms of the Pacific coast of North America, where upwelling-related variables were important predictors of biogeographic structure for algae and invertebrates with lower to medium dispersal capabilities (Fenberg et al., 2014).

### *Variables related to an upwelling explain distributions of the two species in the* R. levigatoides *complex*

In our PCA-env analyses of the two species of the *R. levigatoides* complex, the variables most correlated to the first principal component (Sea Surface Temperature mean, Chlorophyll A mean, and Sea Surface Salinity of the freshest month) were strongly correlated to variables that are indicative to an upwelling, i.e., concentrations of Phosphate, Nitrate and Silicate, and Productivity (Fenberg et al., 2014). Based on environmental niche modeling, in both species of the *R. levigatoides* complex, the most relevant variables were related to currents and upwelling (except by Bathymetry and Minimal light at the bottom), while atmospheric variables, like those related to precipitation and cloud cover, were of small relevance. In contrast, *A. legionium* had a much broader niche, having suitable places scattered along the entire coast. The most relevant factors had almost no differentiation across the Brazilian coast and were related to the intertidal environment, like Bathymetry and Maximal light at the bottom.

#### Temporal context

The two species of the *R. levigatoides* complex diverged about 7.2, 1.69- 16.28 Ma (Appendix 2, Fig. 6), while their COI coalescent analyses recover an older age, 10.01, 3.93-18.5 Ma. Each of the species also had a relatively old COI coalescence times: 5.31, 2.7-8.72 (SE species) and 3.16, 1.62-5.24 (NE species). While older gene coalescence times compared to species divergence times are expected, we note that the divergence of these two cryptic species (Appendix 2, Fig. 5) matches well with the dates of the presumed origin of the Brazilian upwelling, 7.2, 1.69-16.28 vs 7.25-4.8 Ma (see the Introduction section). A similar timing, 7.8, 13.46-3.21 Ma, was detected for *A. bilophus*, which was employed as an outgroup in *A. legionium* analyses (Fig. 5B). COI haplotypes of *A. legionium* had relatively recent coalescence, 2.01, 0.79-3.53 Ma (Fig. 5B). Our estimates of divergence times in the marine mites suggest that the Last Glacial Maximum (LGM, 26,500 years ago) was probably not the main factor shaping the distribution of these organisms. We expect, however, that the lower sea level during the LGM (when AP was dry land) reinforced the isolation between populations of both *R. levigatoides* and *A. legionium*.

#### General considerations

It is remarkable that the characteristic allopatric distributions observed in the NE and SE species of the *R. levigatoides* complex are mirrored by the distributions of two King weakfish sibling species of the genus *Macrodon* (Santos et al., 2006). Similarly, many community level studies (Lotufo, 2002, Floeter et al., 2001, Barroso et al., 2016, Pinheiro et al., 2018) also point to a break northward relative to Cabo Frio. These data indicate the presence of a general biogeographic pattern, most importantly highlighting the Abrolhos Plateau (AP) as a major biogeographic break and the SACW upwelling as major ecological factor affecting niche specialization.

Here we suggest modifying currently practiced sampling strategies to include extensive sampling near the AP region and using more sophisticated models for circulation and environmental parameters. Special care should be taken when planning sampling along the Bahia State coast because its southern portion might be the boundary between tropical and subtropical faunas. Furthermore, our study highlights the importance of the complex current circulation pattern at the Abrolhos Plateau and Trindade Ridge (Peterson & Stramma, 1991), which may act as a physical barrier restricting gene flow between populations. This complex pattern is usually ignored in most marine biogeographic studies (*e.g.* Andrade et al., 2011; Santos et al. 2006).

## Conclusions

Our study supports the hypotheses that the main break in the distribution of coastal rocky-shore communities occurs at Abrolhos Plateau and Vitória-Trindade Ridge, not at Cabo Frio as traditionally accepted in the literature (Briggs, 1974, 1995, Spalding et al, 2007, Petuch, 2013). This region is not only a barrier to gene flow due complex current patterns, but it also is a place where sharp changes in major environmental parameters associated with the South Atlantic Central Waters (SACW) upwelling occur. Our hypothesis predicts parapatric speciation or population structuring with the Abrolhos Plateau limiting the gene flow and the SACW upwelling offering unique ecological niches. Strong empirical evidence supporting our biogeographic hypothesis comes from *Rhombognathus* mites (this study) and King weakfish (*Macrodon*). Finally, the onset of current conditions is relatively old (4.8-7.25 Ma), corresponding to the full activity of the Malvinas (Falklands) current and establishing of SACW.

## ACKNOWLEDGEMENTS

ARP and PBK were supported by Coordenação de Aperfeiçoamento de Pessoal de Nível Superior (CAPES) Ciência sem Fronteiras (Brazil; PVE 88881.064989/2014-01) and the Russian Science Foundation (project No. 19-14-00004). Writing of the final draft of this article was possible due to the support offered to ARP as a postdoc at the Tyumen State University. The authors thank the Program for Technological Development in Tools for Health-PDTISFIOCRUZ for use of its facilities. Special thanks to Pedro Henrique Martins and Arthur Anker for providing samples from Cumuruxatiba (BA) and to Dr. Flávio Fernandes (Instituto de Estudos do Mar Almirante Paulo Moreira) for his support and logistical help for our collecting trip in the Cabo Frio area. Special thanks are to Dr. Barry OConnor for receiving ARP at the University of Michigan (Department of Ecology and Evolutionary Biology) as a visiting researcher in 2018 and reading the final version of the manuscript. Data sets were processed on the Sagarana HPC cluster, CEPAD-ICB-UFMG. Activities involving work with genetic material of Brazilian mites were registered in the SisGen database, accession codes A48017D and A850392.

## BIOSKETCH

Almir Pepato is interested in the use of mites, especially marine mites, as models for biogeography. This work represents the first result from a long term cooperation with the two co-authors in sampling and identifying the mites along the Brazilian coast.

Teofânia Vidigal coordinates the laboratories of Malacology and Molecular Systematics at Zoological Department of UFMG. As part of the investigations concerning the biogeography along the Brazilian coast, she is especially interested in the air breathing marine slugs of the Genus *Onchidella*. She is also studding amphibious slugs and invasive bivalves.

Pavel Klimov has a broad interest on understanding the evolution of Acariformes mites, ranging from genomic evolution, species delimitation, phylogeny and character evolution.

Author contributions: ARP, THDAV and PBK, conceived the project and conducted the fieldwork; ARP was the main responsible for obtaining sequences and analyses; ARP and PBK wrote the paper with the assistance from THDAV.

## Supplementary material

Appendix 1. Population and samples characterization, including details on sampling, sequencing and genetic diversity statistics.

Appendix 2. Timetree inference of acariform mites (Acariformes)

Appendix 3. Mantel test of raw genetic distances vs geographic distances. Left column geographical distances in Km, in right columns transformed in logarithmic (base 10), for *Agauopsis legionium*, the entire *Rhombognathus levigatoides* complex, and separately for *Rhombognathus levigatoides* Northeast and *Rhombognathus levigatoides* Southeast clades..

Appendix 4. Niche comparison and modeling

