## Appendix 1 for "Evaluating the boundaries of marine biogeographic regions of the Southwestern Atlantic using halacarid mites (Halacaridae), meiobenthic organisms with a low dispersal potential"

### Appendix 1. Population and samples characterization

*DNA amplification and sequencing*

Specimens were preserved in 95-100 % ethanol and stored at −20 or −80°C. Genomic DNA was extracted from single specimens using a QIAamp DNA Micro kit (Qiagen) following the manufacturer's protocol except by using two steps of the final elution, leading to a final volume of 70 µl.

The amplification of COI was conducted using nested PCR with parent PCR product being produced employing forward primer COX1_16F (TGANTWTTTTCHACWAAYCAYAA) or alternatively COX1_220F (ATAATHGGDGGDTTYGGIAA) and the reverse primer COX1_1324R (CDGWRTAHCGDCGDGGTAT). The second PCR was performed with primers including M13 forward and reverse tails, respectively (not shown on the sequences): COX1_270F_T (TGAYATRGCNTWYCCICG) COX1 917RT (DGTRAARTADGCHCGDGTRTC). Both rounds of amplifications were performed in 20 µl of final volume, with Platinum Taq DNA Polymerase (Invitrogen) in a Mastercycler gradient, Eppendorf thermocycler. Master mix for initial PCR contained 2.0 µl of PCR buffer (1X), 1.4 µl $MgSO_4$ (50 mM), 1.4 µL of dNTP (10 mM each), 0.8 µl of each oligonucleotide primer. The first round of amplification used 1-3 µl of genomic DNA and included an initial denaturing step of 4 min at 94°C, ten amplification cycles, comprising: denaturing at 94°C for 30s, annealing at 48°C for 35s, and extension at 72°C for 2 min, 18 amplification cycles changing the annealing temperature to 51°C, and a final step of extension of 5 min at 72°C. The second round employed 0.5 µl from the parent PCR product and included an initial denaturing step of 4 min at 94°C, 35 amplification cycles, comprising: denaturing at 94°C for 30s, annealing at 45°C for 60 s, and extension at 72°C for 2 min, and a final step of extension of 5 min. at 72°C. Amplification protocols for other loci are given in Pepato et al. (2018).

PCR products were purified using the Ampure® (Agencourt) kit and sequenced using a 3730 DNA Analyzer. M13 FORW/REV primers were used for sequencing. Chromatograms were resolved in ChromasPro 1.41 (Technelysium Pty Ltd).

### Results

*Agauopsis legionium*

Eighty-nine sequenced individuals formed 11 populations distributed from the Amazonian region of Pará State in the north (0°36'S 47°20'W) to Santa Catarina State in the south (26°48' S 48°35'W) (Table S1, Fig. 1). The total alignment length was 609 nucleotide positions, of which 70 sites were variable and 46 parsimony informative; after amino acid translation, 12 amino acid positions were variable, and six were parsimony informative. The data set presented 58 different haplotypes and a haplotype diversity Hd = 0,9816. The nucleotide diversity, $\pi = 10.95 \pm 5.6$.

*Rhombognathus levigatoides*

One hundred twenty eight sequenced individuals were grouped in 14 populations, ranging from the Parnaíba River Delta, Piaui State to Torres, Rio Grande do Sul State (2° 48' S to 29° 21'S). Out of 609 nucleotide alignment positions, 150 were variable and 132 were parsimony informative; 13 amino acid residues were variable, and nine of them were parsimony informative. There were 99 distinct haplotypes and with Haplotype diversity Hd = 0.9940; the nucleotide diversity was $\pi = 52.21 \pm 25.2$ (Table S1).

**Table S1**. Sequenced individuals of *Rhombognathus levigatoides* Pepato & Rocha, 2007 complex and *Agauopsis legionium* Pepato & Tiago, 2005. Provinces and ecoregions of the Brazilian coast follow Spalding et al. (2007). When applicable, nucleotide (π) and haplotypic (Hd) diversities are given as mean ± SD.

| **Province** | **Ecoregion** | **#Locality** | **Latitude** | **Longitude** | ***R. levigatoides*** | ***A. legionium*** | **Pop.** | ***R. levigatoides*** | ***A. legionium*** |
|---|---|---|---|---|---|---|---|---|---|
| North Brazil Shelf | Amazonia | 1. PA | -0.596325 | -47.330775 | -- | 16 | N01 | NA | π= 0.00321±0.00082; Hd= 0.831± 0.085 |
| | | 2. MA | -2.490997 | -44.306447 | -- | 01 | | | |
| Tropical Southwestern Atlantic | Northeastern Brazil | 3. PI | -2.803239 | -41.729286 | 05 | -- | NE1 | π=0.00493±0.00150; Hd=1.000±0.01600 | NA |
| | | 4. PI | -2.922886 | -41.344211 | -- | 01 | NE2 | π=0.01026±0.00289; Hd=0.972±0.064 | π=0.00239±0.00071; Hd= 0.727±0.144 |
| | | 5. CE | -3.688481 | -38.611197 | 01 | 04 | | | |
| | | **6.** CE | -3.686830 | -38.640300 | 08 | 06 | | | |
| | | 7. RN | -6.226086 | -35.044461 | 09 | 06 | NE3 | π=0.00607±0.00159; Hd=0.944±0.070 | π=0.00372±0.00143; Hd=0.867±0.129 |
| | | 8. PB | -6.686881 | -34.930664 | 08 | -- | NE4 | π=0.02673±0.00749; Hd=0.944±0.070 | NA |
| | | 9. PB | -7.145014 | -34.806472 | 01 | -- | | | |
| | | 10. PE | -7.630539 | -34.808539 | 01 | 07 | NE5 | π=0.00339± 0.00081; Hd=0.867±0.107 | π= 0.01345±0.00419; Hd=0.952±0.096 |
| | | 11. PE | -8.457536 | -34.982994 | 08 | -- | | | |
| | | 12. PE | -8.302775 | -34.947014 | 01 | -- | | | |
| | | 13. AL | -9.840211 | -35.889522 | 12 | -- | NE6 | π=0.00440±0.00095; Hd=0.909±0.079 | NA |
| | Eastern Brazil | 14. BA | -13.00728 | -38.454197 | 02 | -- | E01 | π=0.01943±0.00306; Hd=0.861±0.087 | π=0.00361±0.00090; Hd=0.933±0.122 |
| | | 15. BA | -13.01021 | -38.520697 | 06 | 06 | | | |
| | | 16. BA | -14.28396 | -38.984781 | 04 | 05 | E02 | π=0.01595±0.00266; Hd=1.000±0.076 | π=0.00164±0.00057; Hd=0.700± 0.218 |
| | | 17. BA | -14.92399 | -39.017847 | 03 | -- | | | |
| | | 18. BA | -17.063500 | -39.170389 | 05 | -- | E03 | π=0.01018±0.00181; Hd=1.000±0.126 | NA |
| | | 19. ES | -19.93689 | -40.132667 | -- | 10 | E04 | π=0.00775±0.00122; | π=0.00234±0.00054; |

| | | | | | | | | | |
|---|---|---|---|---|---|---|---|---|---|
| | | 20. ES | -20.031994 | -40.159044 | 01 | -- | | Hd=0.889±0.091 | Hd=0.844±0.103 |
| | | 21. ES | -20.038350 | -40.177894 | 08 | -- | | | |
| Warm Temperate Southwestern Atlantic | Southeastern Brazil | 22. RJ | -22.598858 | -41.990372 | 01 | -- | SE1 | π=0.02716±0.00391; Hd=0.956±0.045 | π=0.00788±0.00150; Hd=0.900±0.161 |
| | | 23. RJ | -22.96143 | -42.020378 | 03 | 01 | | | |
| | | 24. RJ | -22.96414 | -42.012556 | 08 | -- | | | |
| | | 25. RJ | -22.97928 | -42.019367 | 02 | 01 | | | |
| | | 26. RJ | -22.98149 | -42.038006 | -- | 03 | | | |
| | | 27. SP | -23.57601 | -45.308014 | -- | 02 | SE2 | π=0.01861±0.00431; Hd=0.952±0.096 | π= 0.01235±0.00297; Hd=0.905±0.103 |
| | | 28. SP | -23.82197 | -45.410717 | 07 | 09 | | | |
| | | 29. SP | -25.20194 | -47.984406 | -- | 04 | SE3 | NA | π=0.01423± 0.00356; Hd= 1.000± 0.177 |
| | | 30. SC | -26.774389 | -48,636583 | 01 | -- | SE4 | π=0.02015±0.00196; Hd=1.000±0.034 | π= 0.01314±0.00245 Hd=0.905±0.01067 |
| | | 31. SC | -26,780750 | -48,603583 | 04 | -- | | | |
| | | 32. SC | -26.80260 | -48.596133 | -- | 04 | | | |
| | | 33. SC | -26,924150 | -48,634781 | 05 | -- | | | |
| | | 34. SC | -27.74820 | -48.499236 | 01 | 03 | | | |
| | Rio Grande | 35. RS | -29.358381 | -49.733639 | 11 | --- | RG1 | π=0.02365±0.00348; Hd=0.891±0.074 | NA |

**Table S2.** GenBank accession numbers and vouchering of newly sequenced COI for individuals belonging to four morphospecies of the genera *Agauopsis* and *Rhombognathus*.

| Vouchering (UFMG-AC) | Locality | Coordinates (Lat./Long) | GenBank accession |
|---|---|---|---|
| *Agauopsis bilophus* | | | |
| 1301135 | Pedra do Xáreu, Cabo de Santo Agostinho, PE, 10.VIII.2007 | -8.30492778,-34.9482611 | MH999696 |
| 1301136 | Pedra do Xáreu, Cabo de Santo Agostinho, PE, 10.VIII.2007 | -8.30492778,-34.9482611 | MH999697 |
| 1700728 | Ponta de Pedra, Pituba, Salvador, BA, 20/X/2015 | -13.007275,-38.45419722 | MH999698 |
| 1700729 | Ponta de Pedra, Pituba, Salvador, BA, 20/X/2015 | -13.007275,-38.45419722 | MH999699 |
| 1700730 | Ponta de Pedra, Pituba, Salvador, BA, 20/X/2015 | -13.007275,-38.45419722 | MH999700 |
| 1703950 | Praia do Resende, Itacaré, BA, 30/X/2015 | -14.283958, -38.984781 | MH999701 |
| 1704013 | Praia do Resende, Itacaré, BA, 30/X/2015 | -14.283958, -38.984781 | MH999702 |
| 1300477 | Enseada das Garças, Fundão, ES, 10.VIII.2014 | -20.0319944,-40.1590444 | MH999703 |
| 1301111 | Praia do Forno, Arraial do Cabo, RJ, 19.III.2015 | -22.964136, -42.012556 | MH999704 |
| 1301144 | Ilha de Cabo Frio, Arraial do Cabo, RJ, 18.III.2015 | -22.9986889,-42.0031694 | MH999705 |
| 1301145 | Ilha de Cabo Frio, Arraial do Cabo, RJ, 18.III.2015 | -22.9986889,-42.0031694 | MH999706 |
| 1301142 | Ilha de Cabo Frio, Arraial do Cabo, RJ, 18.III.2015 | -22.9986889,-42.0031694 | MH999707 |
| 1301160 | Prainha, Arraial do Cabo, RJ, 16.III.2015 | -22.961428, -42.020378 | MH999709 |
| 1301149 | Praia das Cabeçudas, Itajaí, SC, 06.II.2015 | -26,924150, -48,634781 | MH999708 |
| 1300506 | Praia do Quilombo, Penha, SC, 05.II.2015 | -26.774389, -48,6365833 | MH999710 |
| 1301148 | Praia das Cabeçudas, Itajaí, SC, 06.II.2015 | -26,924150, -48,634781 | MH999711 |
| *Agauopsis legionium* | | | |
| 1300487 | Praia do Farol Velho, Salinópolis, PA, 09.VII. 2014 | -0.596325, -47.330775 | MH999712 |
| 1300488 | Praia do Farol Velho, Salinópolis, PA, 09.VII. 2014 | -0.596325, -47.330775 | MH999713 |
| 1300496 | Praia do Farol Velho, Salinópolis, PA, 09.VII. 2014 | -0.596325, -47.330775 | MH999714 |
| 1300497 | Praia do Farol Velho, Salinópolis, PA, 09.VII. 2014 | -0.596325, -47.330775 | MH999715 |
| 1301128 | Praia do Farol Velho, Salinópolis, PA, 09.VII. 2014 | -0.596325, -47.330775 | MH999716 |
| 1301129 | Praia do Farol Velho, Salinópolis, PA, 09.VII. 2014 | -0.596325, -47.330775 | MH999717 |
| 1301130 | Praia do Farol Velho, Salinópolis, PA, 09.VII. 2014 | -0.596325, -47.330775 | MH999718 |

| | | | |
|---|---|---|---|
| 1301131 | Praia do Farol Velho, Salinópolis, PA, 09.VII. 2014 | -0.596325, -47.330775 | MH999719 |
| 1301133 | Praia do Farol Velho, Salinópolis, PA, 09.VII. 2014 | -0.596325, -47.330775 | MH999720 |
| 1301162 | Praia do Farol Velho, Salinópolis, PA, 09.VII. 2014 | -0.596325, -47.330775 | MH999721 |
| 1301163 | Praia do Farol Velho, Salinópolis, PA, 09.VII. 2014 | -0.596325, -47.330775 | MH999722 |
| 1301164 | Praia do Farol Velho, Salinópolis, PA, 09.VII. 2014 | -0.596325, -47.330775 | MH999723 |
| 1301165 | Praia do Farol Velho, Salinópolis, PA, 09.VII. 2014 | -0.596325, -47.330775 | MH999724 |
| 1301166 | Praia do Farol Velho, Salinópolis, PA, 09.VII. 2014 | -0.596325, -47.330775 | MH999725 |
| 1301167 | Praia do Farol Velho, Salinópolis, PA, 09.VII. 2014 | -0.596325, -47.330775 | MH999726 |
| 1301168 | Praia do Farol Velho, Salinópolis, PA, 09.VII. 2014 | -0.596325, -47.330775 | MH999727 |
| 1704159 | Ponta de Areia, São Luis, MA, 17.IX.2016 | -2.490997, -44.306447 | MH999728 |
| 1704337 | Cajueiro da Praia, PI, 20.IX.2016 | -2.922886, -41.344211 | MH999729 |
| 1700740 | Praia de Iparaná, Caucaia, CE, 22.IX.2016 | -3.688481, -38.611197 | MH999730 |
| 1700742 | Praia de Iparaná, Caucaia, CE, 22.IX.2016 | -3.688481, -38.611197 | MH999731 |
| 1700745 | Praia de Iparaná, Caucaia, CE, 22.IX.2016 | -3.688481, -38.611197 | MH999732 |
| 1700749 | Praia de Iparaná, Caucaia, CE, 22.IX.2016 | -3.688481, -38.611197 | MH999733 |
| 1704269 | Praia do Pacheco, Caucaia, CE, 21.IX.2016 | -3.686830, -38.640300 | MH999734 |
| 1704270 | Praia do Pacheco, Caucaia, CE, 21.IX.2016 | -3.686830, -38.640300 | MH999735 |
| 1704271 | Praia do Pacheco, Caucaia, CE, 21.IX.2016 | -3.686830, -38.640300 | MH999736 |
| 1704272 | Praia do Pacheco, Caucaia, CE, 21.IX.2016 | -3.686830, -38.640300 | MH999737 |
| 1704273 | Praia do Pacheco, Caucaia, CE, 21.IX.2016 | -3.686830, -38.640300 | MH999738 |
| 1704275 | Praia do Pacheco, Caucaia, CE, 21.IX.2016 | -3.686830, -38.640300 | MH999739 |
| 174039 | Praia de Pipa, Tibau do Sul, RN, 17.II.2017 | -6.226086, -35.044461 | MH999740 |
| 174172 | Praia de Pipa, Tibau do Sul, RN, 17.II.2017 | -6.226086, -35.044461 | MH999745 |
| 174184 | Praia de Pipa, Tibau do Sul, RN, 17.II.2017 | -6.226086, -35.044461 | MH999741 |
| 174185 | Praia de Pipa, Tibau do Sul, RN, 17.II.2017 | -6.226086, -35.044461 | MH999742 |
| 174236 | Praia de Pipa, Tibau do Sul, RN, 17.II.2017 | -6.226086, -35.044461 | MH999743 |
| 174237 | Praia de Pipa, Tibau do Sul, RN, 17.II.2017 | -6.226086, -35.044461 | MH999744 |

| | | | |
|---|---|---|---|
| 174031 | Ponta de Pedra, Goiana, PE, 13.II.2017 | -7.630539, -34.808539 | MH999746 |
| 174032 | Ponta de Pedra, Goiana, PE, 13.II.2017 | -7.630539, -34.808539 | MH999747 |
| 1704033 | Ponta de Pedra, Goiana, PE, 13.II.2017 | -7.630539, -34.808539 | MH999748 |
| 1704180 | Ponta de Pedra, Goiana, PE, 13.II.2017 | -7.630539, -34.808539 | MH999749 |
| 1704182 | Ponta de Pedra, Goiana, PE, 13.II.2017 | -7.630539, -34.808539 | MH999750 |
| 1704183 | Ponta de Pedra, Goiana, PE, 13.II.2017 | -7.630539, -34.808539 | MH999751 |
| 1704190 | Ponta de Pedra, Goiana, PE, 13.II.2017 | -7.630539, -34.808539 | MH999752 |
| 1702586 | Barra, Salvador, BA, 16.VIII.2016 | -13.010208, -38.520697 | MH999753 |
| 1702587 | Barra, Salvador, BA, 16.VIII.2016 | -13.010208, -38.520697 | MH999754 |
| 1702588 | Barra, Salvador, BA, 16.VIII.2016 | -13.010208, -38.520697 | MH999755 |
| 1704199 | Barra, Salvador, BA, 16.VIII.2016 | -13.010208, -38.520697 | MH999756 |
| 1704200 | Barra, Salvador, BA, 16.VIII.2016 | -13.010208, -38.520697 | MH999757 |
| 1704201 | Barra, Salvador, BA, 16.VIII.2016 | -13.010208, -38.520697 | MH999758 |
| 1700739 | Praia do Resende, Itacaré BA, 30.X.2015 | -14.283958, -38.984781 | MH999759 |
| 1704018 | Praia do Resende, Itacaré BA, 30.X.2015 | -14.283958, -38.984781 | MH999760 |
| 1704019 | Praia do Resende, Itacaré BA, 30.X.2015 | -14.283958, -38.984781 | MH999761 |
| 1704020 | Praia do Resende, Itacaré BA, 30.X.2015 | -14.283958, -38.984781 | MH999762 |
| 1704021 | Praia do Resende, Itacaré BA, 30.X.2015 | -14.283958, -38.984781 | MH999763 |
| 174001 | Praia do Padre, Aracruz, ES, 08.X.2014 | -19.936889, -40.132667 | MH999764 |
| 174002 | Praia do Padre, Aracruz, ES, 08.X.2014 | -19.936889, -40.132667 | MH999765 |
| 174003 | Praia do Padre, Aracruz, ES, 08.X.2014 | -19.936889, -40.132667 | MH999766 |
| 174004 | Praia do Padre, Aracruz, ES, 08.X.2014 | -19.936889, -40.132667 | MH999767 |
| 174005 | Praia do Padre, Aracruz, ES, 08.X.2014 | -19.936889, -40.132667 | MH999768 |
| 174006 | Praia do Padre, Aracruz, ES, 08.X.2014 | -19.936889, -40.132667 | MH999769 |
| 174223 | Praia do Padre, Aracruz, ES, 08.X.2014 | -19.936889, -40.132667 | MH999770 |
| 174224 | Praia do Padre, Aracruz, ES, 08.X.2014 | -19.936889, -40.132667 | MH999771 |
| 174225 | Praia do Padre, Aracruz, ES, 08.X.2014 | -19.936889, -40.132667 | MH999772 |

| | | | |
|---|---|---|---|
| 174226 | Praia do Padre, Aracruz, ES, 08.X.2014 | -19.936889, -40.132667 | MH999773 |
| 1301154 | Ilha dos Franceses, Arraial do Cabo, RJ, 19.III.2015 | -22.981494, -42.038006 | MH999774 |
| 1301159 | Praia dos Anjos, Arraial do Cabo, RJ, 17.III.2015 | -22.979281, -42.019367 | MH999775 |
| 1301161 | Prainha, Arraial do Cabo, RJ, 16.III.2015 | -22.961428, -42.020378 | MH999776 |
| 1704384 | Ilha dos Franceses, Arraial do Cabo, RJ, 19.III.2015 | -22.981494, -42.038006 | MH999777 |
| 1704385 | Ilha dos Franceses, Arraial do Cabo, RJ, 19.III.2015 | -22.981494, -42.038006 | MH999778 |
| 174366 | Praia de Mococa, Caraguatatuba, SP, 07.III.2016 | -23.576014, -45.308014 | MH999779 |
| 174367 | Praia de Mococa, Caraguatatuba, SP, 07.III.2016 | -23.576014, -45.308014 | MH999780 |
| 174386 | Praia Preta, São Sebastião, SP, 09.III.2016 | -23.821972, -45.410717 | MH999781 |
| 174387 | Praia Preta, São Sebastião, SP, 09.III.2016 | -23.821972, -45.410717 | MH999782 |
| 174389 | Praia Preta, São Sebastião, SP, 09.III.2016 | -23.821972, -45.410717 | MH999783 |
| 174390 | Praia Preta, São Sebastião, SP, 09.III.2016 | -23.821972, -45.410717 | MH999784 |
| 174391 | Praia Preta, São Sebastião, SP, 09.III.2016 | -23.821972, -45.410717 | MH999785 |
| 174277 | Praia Preta, São Sebastião, SP, 09.III.2016 | -23.821972, -45.410717 | MH999786 |
| 174278 | Praia Preta, São Sebastião, SP, 09.III.2016 | -23.821972, -45.410717 | MH999787 |
| 174279 | Praia Preta, São Sebastião, SP, 09.III.2016 | -23.821972, -45.410717 | MH999788 |
| 174280 | Praia Preta, São Sebastião, SP, 09.III.2016 | -23.821972, -45.410717 | MH999789 |
| 1301018 | Marujá, Ilha do Cardoso, Cananeia, SP, 02.I.2012 | -25.201944, -47.984406 | MH999790 |
| 1301151 | Marujá, Ilha do Cardoso, Cananeia, SP, 02.I.2012 | -25.201944, -47.984406 | MH999791 |
| 1301152 | Marujá, Ilha do Cardoso, Cananeia, SP, 02.I.2012 | -25.201944, -47.984406 | MH999792 |
| 1301153 | Marujá, Ilha do Cardoso, Cananeia, SP, 02.I.2012 | -25.201944, -47.984406 | MH999793 |
| 1301155 | Praia Vermelha, Penha, SC, 05.II.2015 | -26.802603, -48.596133 | MH999794 |
| 1301156 | Praia Vermelha, Penha, SC, 05.II.2015 | -26.802603, -48.596133 | MH999795 |
| 1301157 | Praia Vermelha, Penha, SC, 05.II.2015 | -26.802603, -48.596133 | MH999796 |
| 1301158 | Praia Vermelha, Penha, SC, 05.II.2015 | -26.802603, -48.596133 | MH999797 |
| 174381 | Praia da Armação, Florianópolis, SC, 07.II.2015 | -27.748200, -48.499236 | MH999798 |
| 174382 | Praia da Armação, Florianópolis, SC, 07.II.2015 | -27.748200, -48.499236 | MH999799 |

| | | | |
|---|---|---|---|
| 174383 | Praia da Armação, Florianópolis, SC, 07.II.2015 | -27.748200, -48.499236 | MH999800 |
| *Rhombognathus areolatus* | | | |
| 1704174 | Praia das Trincheiras, Baía da Traição, PB, 16/II/2017 | -6.686881,-34.930664 | MH999551 |
| 1704008 | Pedra do Xáreu, Itapoama, PE, 12/II/2017 | -8.302775, -34.947014 | MH999552 |
| 1704178 | Pedra do Xáreu, Itapoama, PE, 12/II/2017 | -8.302775, -34.947014 | MH999553 |
| 1704192 | Praia da Sereia, Maceió, AL, 21/VIII/2016 | -9.566364,-35.645336 | MH999554 |
| 1704193 | Praia da Sereia, Maceió, AL, 21/VIII/2016 | -9.566364,-35.645336 | MH999555 |
| 1703991 | Ponta de Pedra, Pituba, Salvador, BA, 20/X/2015 | -13.007275,-38.454197 | MH999556 |
| 1704197 | Praia do Moreira, Cumuruxatiba, Prado, BA, 5/I/2015 | -17.0635,-39.170389 | MH999557 |
| 1704196A | Praia do Moreira, Cumuruxatiba, Prado, BA, 5/I/2015 | -17.0635,-39.170389 | MH999558 |
| 1704195 | Praia do Moreira, Cumuruxatiba, Prado, BA, 5/I/2015 | -17.0635,-39.170389 | MH999559 |
| 1704196B | Praia do Moreira, Cumuruxatiba, Prado, BA, 5/I/2015 | -17.0635,-39.170389 | MH999560 |
| 1300473 | Enseada das Garças, Fundão, ES, 10.VIII.2014 | -20.0319944,-40.1590444 | MH999561 |
| 1600622 | Enseada das Garças, Fundão, ES, 10.VIII.2014 | -20.0319944,-40.1590444 | MH999562 |
| 1600623 | Enseada das Garças, Fundão, ES, 10.VIII.2014 | -20.0319944,-40.1590444 | MH999563 |
| 1600625 | Enseada das Garças, Fundão, ES, 10.VIII.2014 | -20.0319944,-40.1590444 | MH999564 |
| 1600643 | Enseada das Garças, Fundão, ES, 10.VIII.2014 | -20.0319944,-40.1590444 | MH999565 |
| 1600646 | Enseada das Garças, Fundão, ES, 10.VIII.2014 | -20.0319944,-40.1590444 | MH999566 |
| 1600647 | Enseada das Garças, Fundão, ES, 10.VIII.2014 | -20.0319944,-40.1590444 | MH999567 |
| *Rhombognathus levigatoides* | | | |
| 172573 | Pedra do Sal, Parnaíba, PI, 19.IX.2016 | -2.03853, -41.730247 | MH999569 |
| 174400 | Pedra do Sal, Parnaíba, PI, 19.IX.2016 | -2.03853, -41.730247 | MH999568 |
| 174411 | Pedra do Sal, Parnaíba, PI, 19.IX.2016 | -2.03853, -41.730247 | MH999570 |
| 174304 | Pedra do Sal, Parnaíba, PI, 19.IX.2016 | -2.03853, -41.730247 | MH999571 |
| 174362 | Pedra do Sal, Parnaíba, PI, | -2.03853, - | MH999572 |

| | | | |
|---|---|---|---|
| | 19.IX.2016 | 41.730247 | |
| 170750 | Praia de Iparaná, Caucaia, CE, 22.IX.2016 | -2.03853, - 41.730247 | MH999573 |
| 1704058 | Praia do Pacheco, Caucaia, CE, 21.IX.2016 | -3.686830, - 38.640300 | MH999574 |
| 1704221 | Praia do Pacheco, Caucaia, CE, 21.IX.2016 | -3.686830, - 38.640300 | MH999575 |
| 1704369A | Praia do Pacheco, Caucaia, CE, 21.IX.2016 | -3.686830, - 38.640300 | MH999576 |
| 1704369B | Praia do Pacheco, Caucaia, CE, 21.IX.2016 | -3.686830, - 38.640300 | MH999577 |
| 1704369C | Praia do Pacheco, Caucaia, CE, 21.IX.2016 | -3.686830, - 38.640300 | MH999578 |
| 1704265 | Praia do Pacheco, Caucaia, CE, 21.IX.2016 | -3.686830, - 38.640300 | MH999579 |
| 1704370 | Praia do Pacheco, Caucaia, CE, 21.IX.2016 | -3.686830, - 38.640300 | MH999580 |
| 1704267 | Praia do Pacheco, Caucaia, CE, 21.IX.2016 | -3.686830, - 38.640300 | MH999581 |
| 1704242 | Praia de Pipa, Tibau do Sul, RN, 17.II.2017 | -6.226086, - 35.044461 | MH999582 |
| 1704243 | Praia de Pipa, Tibau do Sul, RN, 17.II.2017 | -6.226086, - 35.044461 | MH999583 |
| 1704244 | Praia de Pipa, Tibau do Sul, RN, 17.II.2017 | -6.226086, - 35.044461 | MH999584 |
| 1704295 | Praia de Pipa, Tibau do Sul, RN, 17.II.2017 | -6.226086, - 35.044461 | MH999585 |
| 1704297 | Praia de Pipa, Tibau do Sul, RN, 17.II.2017 | -6.226086, - 35.044461 | MH999586 |
| 1704405 | Praia de Pipa, Tibau do Sul, RN, 17.II.2017 | -6.226086, - 35.044461 | MH999587 |
| 1704409 | Praia de Pipa, Tibau do Sul, RN, 17.II.2017 | -6.226086, - 35.044461 | MH999588 |
| 1704410 | Praia de Pipa, Tibau do Sul, RN, 17.II.2017 | -6.226086, - 35.044461 | MH999589 |
| 1704301 | Praia de Pipa, Tibau do Sul, RN, 17.II.2017 | -6.226086, - 35.044461 | MH999590 |
| 1704255 | Arrecife do Farol, Baia da Traição, PB, 16.II.2017 | -6.686881, - 34.930664 | MH999591 |
| 1704365 | Arrecife do Farol, Baia da Traição, PB, 16.II.2017 | -6.686881, - 34.930664 | MH999592 |
| 1704256 | Arrecife do Farol, Baia da Traição, PB, 16.II.2017 | -6.686881, - 34.930664 | MH999593 |
| 1704309 | Arrecife do Farol, Baia da Traição, PB, 16.II.2017 | -6.686881, - 34,930664 | MH999594 |
| 1704310 | Arrecife do Farol, Baia da Traição, PB, 16.II.2017 | -6.686881, - 34.930664 | MH999595 |
| 1704313 | Arrecife do Farol, Baia da Traição, PB, 16.II.2017 | -6.686881, - 34.930664 | MH999596 |
| 1704363 | Arrecife do Farol, Baia da | -6.686881, - | MH999597 |

| | | | |
|---|---|---|---|
| | Traição, PB, 16.II.2017 | 34.930664 | |
| 1704315 | Arrecife do Farol, Baia da Traição, PB, 16.II.2017 | -6.686881, - 34.930664 | MH999598 |
| 1704246 | Cabo Branco, João Pessoa, PB, 15.II.2017 | -7.145014, - 34.806472 | MH999599 |
| 1703982 | Pontal do Cupe, Ipojuca, PE, 12.II.2017 | -8.457536, - 39.982994 | MH999601 |
| 1703984 | Pontal do Cupe, Ipojuca, PE, 12.II.2017 | -8.457536, - 39.982994 | MH999602 |
| 1703985 | Pontal do Cupe, Ipojuca, PE, 12.II.2017 | -8.457536, - 39.982994 | MH999603 |
| 1704012 | Pedra do Xáreu, Cabo de Santo Agostinho, PE, 12.II.2017 | -8.302775, - 34.947014 | MH999604 |
| 1704023 | Ponta de Pedra, Goiana, PE, 13. II. 2017 | -7.629164, - 34.808539 | MH999600 |
| 1704393A | Pontal do Cupe, Ipojuca, PE, 12.II.2017 | -8.457536, - 39.982994 | MH999605 |
| 1704393B | Pontal do Cupe, Ipojuca, PE, 12.II.2017 | -8.457536, - 39.982994 | MH999606 |
| 1704300A | Pontal do Cupe, Ipojuca, PE, 12.II.2017 | -8.457536, - 39.982994 | MH999607 |
| 1704300B | Pontal do Cupe, Ipojuca, PE, 12.II.2017 | -8.457536, - 39.982994 | MH999608 |
| 1704394 | Pontal do Cupe, Ipojuca, PE, 12.II.2017 | -8.457536, - 39.982994 | MH999609 |
| 1704049 | Barra de São Miguel, Maceió, AL, 20.VIII.2016 | -9.840211, - 35.889522 | MH999610 |
| 1704258 | Barra de São Miguel, Maceió, AL, 20.VIII.2016 | -9.840211, - 35.889522 | MH999611 |
| 1704259 | Barra de São Miguel, Maceió, AL, 20.VIII.2016 | -9.840211, - 35.889522 | MH999612 |
| 1704286 | Barra de São Miguel, Maceió, AL, 20.VIII.2016 | -9.840211, - 35.889522 | MH999617 |
| 1704287 | Barra de São Miguel, Maceió, AL, 20.VIII.2016 | -9.840211, - 35.889522 | MH999618 |
| 1704290 | Barra de São Miguel, Maceió, AL, 20.VIII.2016 | -9.840211, - 35.889522 | MH999619 |
| 1704291 | Barra de São Miguel, Maceió, AL, 20.VIII.2016 | -9.840211, - 35.889522 | MH999620 |
| 1704294 | Barra de São Miguel, Maceió, AL, 20.VIII.2016 | -9.840211, - 35.889522 | MH999621 |
| 1704305 | Barra de São Miguel, Maceió, AL, 20.VIII.2016 | -9.840211, - 35.889522 | MH999613 |
| 1704414 | Barra de São Miguel, Maceió, AL, 20.VIII.2016 | -9.840211, - 35.889522 | MH999614 |
| 1704415 | Barra de São Miguel, Maceió, AL, 20.VIII.2016 | -9.840211, - 35.889522 | MH999615 |
| 1704306 | Barra de São Miguel, Maceió, AL, 20.VIII.2016 | -9.840211, - 35.889522 | MH999616 |
| 1703990 | Pituba Ponta de Pedra, Salvador, BA, 28.X.2015 | -13.007275, - 38,454197 | MH999622 |
| 1703993 | Pituba Ponta de Pedra, Salvador, BA, 28.X.2015 | -13.007275, - 38,454197 | MH999623 |
| 1704217A | Barra, Salvador, BA, | -13.010208, - | MH999624 |

| | | | |
|---|---|---|---|
| | 16.VIII.2016 | 38.520697 | |
| 1704217B | Barra, Salvador, BA, 16.VIII.2016 | -13.010208, -38.520697 | MH999625 |
| 1704218 | Barra, Salvador, BA, 16.VIII.2016 | -13.010208, -38.520697 | MH999626 |
| 1704374 | Barra, Salvador, BA, 16.VIII.2016 | -13.010208, -38.520697 | MH999627 |
| 1704375 | Barra, Salvador, BA, 16.VIII.2016 | -13.010208, -38.520697 | MH999628 |
| 1704376 | Barra, Salvador, BA, 16.VIII.2016 | -13.010208, -38.520697 | MH999629 |
| 1704318 | Barra, Salvador, BA, 16.VIII.2016 | -13.010208, -38.520697 | MH999630 |
| 1600666 | Praia do Resende, Itacaré, BA, 30.X.2015 | -14.283958, -38.984781 | MH999631 |
| 1600668 | Praia do Resende, Itacaré, BA, 30.X.2015 | -14.283958, -38.984781 | MH999632 |
| 1600677 | Praia de Olivença, Olivença, BA, 31.X.2015 | -14.923994, -39.017847 | MH999633 |
| 1600678 | Praia de Olivença, Olivença, BA, 31.X.2015 | -14.923994, -39,017847 | MH999634 |
| 1600680 | Praia de Olivença, Olivença, BA, 31.X.2015 | -14.923994, -39,017847 | MH999635 |
| 1704230 | Praia do Resende, Itacaré, BA, 30.X.2015 | -14.283958, -38.984781 | MH999636 |
| 1704231 | Praia do Resende, Itacaré, BA, 30.X.2015 | -14.283958, -38.984781 | MH999637 |
| 1704198A | Praia do Moreira, Cumuruxatiba, Prado, BA, 5.I.2015 | -17.063500, -39.170389 | MH999638 |
| 1704198B | Praia do Moreira, Cumuruxatiba, Prado, BA, 5.I.2015 | -17.063500, -39.170389 | MH999639 |
| 1704234A | Praia do Moreira, Cumuruxatiba, Prado, BA, 5.I.2015 | -17.063500, -39.170389 | MH999640 |
| 1704234B | Praia do Moreira, Cumuruxatiba, Prado, BA, 5.I.2015 | -17.063500, -39.170389 | MH999641 |
| 1704235 | Praia do Moreira, Cumuruxatiba, Prado, BA, 5.I.2015 | -17.063500, -39.170389 | MH999642 |
| 1300479 | Enseada das Garças, Fundão,ES, 10.VIII.2014 | -20.031994, -40,159044 | MH999643 |
| 1704392 | Praia Grande, Fundão, ES, 06.X.2014 | -20.038350, -40.177894 | MH999644 |
| 1704327A | Praia Grande, Fundão, ES, 06.X.2014 | -20.038350, -40.177894 | MH999645 |
| 1704327B | Praia Grande, Fundão, ES, 06.X.2014 | -20.038350, -40.177894 | MH999646 |
| 1704328 | Praia Grande, Fundão, ES, 06.X.2014 | -20.038350, -40.177894 | MH999647 |
| 1704329 | Praia Grande, Fundão, ES, 06.X.2014 | -20.038350, -40.177894 | MH999648 |

| | | | |
|---|---|---|---|
| 1704355 | Praia Grande, Fundão, ES, 06.X.2014 | -20.038350, -40.177894 | MH999649 |
| 1704356 | Praia Grande, Fundão, ES, 06.X.2014 | -20.038350, -40.177894 | MH999650 |
| 1704357 | Praia Grande, Fundão, ES, 06.X.2014 | -20.038350, -40.177894 | MH9996511 |
| 1301026 | Barra de São João, Casimiro de Abreu, RJ, | -22.598858, -41.990372 | MH999652 |
| 1301114 | Praia do Forno, Arraial do Cabo, RJ, 19.III.2015 | -22.964136, -42.012556 | MH999653 |
| 1301115 | Praia do Forno, Arraial do Cabo, RJ, 19.III.2015 | -22.964136, -42.012556 | MH999654 |
| 1600635 | Praia do Forno, Arraial do Cabo, RJ, 19.III.2015 | -22.964136, -42.012556 | MH999655 |
| 1600637 | Praia do Forno, Arraial do Cabo, RJ, 19.III.2015 | -22.964136, -42.012556 | MH999656 |
| 1600638 | Praia do Forno, Arraial do Cabo, RJ, 19.III.2015 | -22.964136, -42.012556 | MH999657 |
| 1600639 | Praia do Forno, Arraial do Cabo, RJ, 19.III.2015 | -22.964136, -42.012556 | MH999658 |
| 1600640 | Praia do Forno, Arraial do Cabo, RJ, 19.III.2015 | -22.964136, -42.012556 | MH999659 |
| 1600654 | Praia dos Anjos, Arraial do Cabo, RJ, 17.III.2015 | -22.979344, -42,019681 | MH999660 |
| 1600655 | Praia dos Anjos, Arraial do Cabo, RJ, 17.III.2015 | -22.979344, -42,019681 | MH999661 |
| 1600661 | Praia do Forno, Arraial do Cabo, RJ, 19.III.2015 | -22.964136, -42.012556 | MH999662 |
| 1704227 | Prainha, Arraial do Cabo, RJ, 16.III.2015 | -22.961711, -42,020800 | MH999663 |
| 174228A | Prainha, Arraial do Cabo, RJ, 16.III.2015 | -22.961711, -42,020800 | MH999664 |
| 174228B | Prainha, Arraial do Cabo, RJ, 16.III.2015 | -22.961711, -42.020800 | MH999665 |
| 1600620 | Praia Preta, São Sebastião, SP, 09.III.2016 | -23.821972, -45.410717 | MH999666 |
| 1700734 | Praia das Pitangueiras, São Sebastião, SP, 09.III.2016 | -23.821972, -45.410717 | MH999667 |
| 1700735 | Praia das Pitangueiras, São Sebastião, SP, 09.III.2016 | -23.821972, -45.410717 | MH999668 |
| 1700736 | Praia das Pitangueiras, São Sebastião, SP, 09.III.2016 | -23.821972, -45.410717 | MH999669 |
| 1700737 | Praia das Pitangueiras, São Sebastião, SP, 09.III.2016 | -23.821972, -45.410717 | MH999670 |
| 1704321 | Praia das Pitangueiras, São Sebastião, SP, 09.III.2016 | -23.821972, -45.410717 | MH999671 |
| 1704323 | Praia das Pitangueiras, São Sebastião, SP, 09.III.2016 | -23.821972, -45.410717 | MH999672 |
| 1300507 | Praia do Quilombo, Penha, SC, 05.II.2015 | -26.774389, -48,6365833 | MH999673 |
| 1600594 | Praia das Cabeçudas, Itajaí, SC, 06.II.2015 | -26,924150, -48,634781 | MH999674 |
| 1600595 | Praia das Cabeçudas, Itajaí, SC, 06.II.2015 | -26,924150, -48,634781 | MH999675 |

| | | | |
|---|---|---|---|
| 1600596 | Praia das Cabeçudas, Itajaí, SC, 06.II.2015 | -26,924150, -48,634781 | MH999676 |
| 1600597 | Praia das Cabeçudas, Itajaí, SC, 06.II.2015 | -26,924150, -48,634781 | MH999677 |
| 1600598 | Praia das Cabeçudas, Itajaí, SC, 06.II.2015 | -26,924150, -48,634781 | MH999678 |
| 1600599 | Praia das Cabeçudas, Itajaí, SC, 06.II.2015 | -26,924150, -48,634781 | MH999679 |
| 1600601 | Praia da Armação, Armação, SC, 05.II.2017 | -26,780750, -48,603583 | MH999680 |
| 1600602 | Praia da Armação, Armação, SC, 05.II.2017 | -26,780750, -48,603583 | MH999681 |
| 1600603 | Praia da Armação, Armação, SC, 05.II.2017 | -26,780750, -48,603583 | MH999682 |
| 1600605 | Praia da Armação, Armação, SC, 05.II.2017 | -26,780750, -48,603583 | MH999683 |
| 1600606 | Praia da Armação, Florianópolis, SC, 07.II.2015 | -27,750419, -48,499997 | MH999684 |
| 1301096 | Parque da Guarita, Torres, RS, 03.II.2015 | -29.358381, -49.733639 | MH999685 |
| 1301097 | Parque da Guarita, Torres, RS, 03.II.2015 | -29.358381, -49.733639 | MH999686 |
| 1301098 | Parque da Guarita, Torres, RS, 03.II.2015 | -29.358381, -49.733639 | MH999687 |
| 1301099 | Parque da Guarita, Torres, RS, 03.II.2015 | -29.358381, -49.733639 | MH999688 |
| 1600586 | Parque da Guarita, Torres, RS, 03.II.2015 | -29.358381, -49.733639 | MH999689 |
| 1600587 | Parque da Guarita, Torres, RS, 03.II.2015 | -29.358381, -49.733639 | MH999690 |
| 1600591 | Parque da Guarita, Torres, RS, 03.II.2015 | -29.358381, -49.733639 | MH999691 |
| 1600610 | Parque da Guarita, Torres, RS, 03.II.2015 | -29.358381, -49.733639 | MH999692 |
| 1600612 | Parque da Guarita, Torres, RS, 03.II.2015 | -29.358381, -49.733639 | MH999693 |
| 1600613 | Parque da Guarita, Torres, RS, 03.II.2015 | -29.358381, -49.733639 | MH999694 |
| 1600614 | Parque da Guarita, Torres, RS, 03.II.2015 | -29.358381, -49.733639 | MH999695 |

**Table S3**. Φ-st statistic calculated using the Tamura and Nei method for *Agauopsis legionium* species complex. Non-significant Φ-st are in bold (significance level=0.05).

| | N01 | NE2 | NE3 | NE5 | E01 | E02 | E04 | SE1 | SE2 | SE3 | SE4 |
|---|---|---|---|---|---|---|---|---|---|---|---|
| N01 | 0.00000 | | | | | | | | | | |
| NE2 | 0.89242 | 0.00000 | | | | | | | | | |
| NE3 | 0.88192 | 0.59907 | 0.00000 | | | | | | | | |
| NE5 | 0.71481 | 0.25331 | 0.26809 | 0.00000 | | | | | | | |
| E01 | 0.87225 | 0.86887 | 0.83775 | 0.51543 | 0.00000 | | | | | | |
| E02 | 0.87576 | 0.88705 | 0.86226 | 0.48914 | 0.44859 | 0.00000 | | | | | |
| E04 | 0.85452 | 0.87685 | 0.85956 | 0.56794 | 0.81052 | 0.83282 | 0.00000 | | | | |
| SE1 | 0.80991 | 0.81883 | 0.76620 | 0.45589 | 0.69987 | 0.70381 | 0.18103 | 0.00000 | | | |
| SE2 | 0.73295 | 0.72201 | 0.67598 | 0.42266 | 0.57114 | 0.53937 | 0.18611 | **0.05790** | 0.00000 | | |
| SE3 | 0.78464 | 0.65316 | 0.57857 | 0.19198 | 0.48704 | 0.41666 | 0.50931 | 0.33934 | 0.26973 | 0.00000 | |
| SE4 | 0.74800 | 0.71309 | 0.64779 | 0.38237 | 0.51065 | 0.44117 | 0.40835 | **0.21403** | **0.08646** | **0.07190** | 0.00000 |

**Table S4**. Φ-st statistic calculated using the Tamura and Nei method for *Rhombognthus levigatoides* populations. Non-significant Φ-st are in bold (significance level=0.05)

| | NE1 | NE2 | NE3 | NE4 | NE5 | NE6 | E01 | E02 | E03 | E04 | SE1 | SE2 | SE3 | RG1 |
|---|---|---|---|---|---|---|---|---|---|---|---|---|---|---|
| NE1 | 0.00000 | | | | | | | | | | | | | |
| NE2 | 0.40460 | 0.00000 | | | | | | | | | | | | |
| NE3 | 0.57463 | **0.02715** | 0.00000 | | | | | | | | | | | |
| NE4 | **0.20577** | **0.08556** | 0.13075 | 0.00000 | | | | | | | | | | |
| NE5 | 0.63013 | 0.51697 | 0.64493 | 0.25152 | 0.00000 | | | | | | | | | |
| NE6 | 0.62503 | 0.54297 | 0.65019 | 0.28886 | 0.35673 | 0.00000 | | | | | | | | |
| E01 | 0.76401 | 0.75249 | 0.78559 | 0.54037 | 0.80718 | 0.80612 | 0.00000 | | | | | | | |
| E02 | 0.81877 | 0.79080 | 0.83034 | 0.56803 | 0.86066 | 0.85579 | 0.31417 | 0.00000 | | | | | | |
| E03 | 0.94525 | 0.92481 | 0.94515 | 0.83910 | 0.95740 | 0.95398 | 0.87504 | 0.89870 | 0.00000 | | | | | |
| E04 | 0.95313 | 0.93674 | 0.95100 | 0.87278 | 0.96023 | 0.95748 | 0.89356 | 0.91716 | 0.88864 | 0.00000 | | | | |
| SE1 | 0.84633 | 0.84956 | 0.85973 | 0.80207 | 0.86550 | 0.86953 | 0.82202 | 0.83188 | 0.72434 | 0.54556 | 0.00000 | | | |
| SE2 | 0.90420 | 0.89682 | 0.91370 | 0.82272 | 0.92587 | 0.92625 | 0.85826 | 0.87503 | 0.81984 | 0.75014 | 0.13997 | 0.00000 | | |
| SE3 | 0.88304 | 0.88113 | 0.89339 | 0.82473 | 0.90151 | 0.90358 | 0.85086 | 0.86353 | 0.78336 | 0.67177 | 0.08675 | 0.14125 | 0.00000 | |
| RG4 | 0.87079 | 0.87140 | 0.88394 | 0.81367 | 0.89269 | 0.89602 | 0.83433 | 0.84951 | 0.75722 | 0.56704 | 0.09289 | 0.24089 | 0.16015 | 0.00000 |
