## Appendix 2 for "Evaluating the boundaries of marine biogeographic regions of the Southwestern Atlantic using halacarid mites (Halacaridae), meiobenthic organisms with a low dispersal potential"

### Appendix 1: Timetree inference of acariform mites (Acariformes)

Since paleontological data for marine mites are absent, we conducted a phylogenetic analysis using outgroup time calibration. A similar approach was used for feather mites resulting in time estimates largely compatible with independent dating of the avian host biogeographic events (Klimov *et al.*, 2017). Our outgroup time calibration analysis was based on previous works (Pepato & Klimov, 2015, Klimov et al., 2018, Dabert et al, 2016, Pepato et al., 2018), but we included a representative set of halacarid marine mite terminals, with all our target species. Species divergence time estimates obtained by this analysis were then used to inform the inference of coalescence times for Cytochrome Oxidase I for our species/population-level datasets.

### Material and Methods

*Sampling and Sequencing*. Taxon names, taxonomic classification, sequenced loci (COI, 18S, 28S) and their GenBank accession numbers are listed in Table S1. For molecular work, we used previously described amplification and sequencing protocols and primers (Klimov et al., 2018, Pepato et al., 2018).

*Molecular analyses*. Mitochondrial COI alignment was unambiguous (no indels); stop-codons, which are indicative of pseudogenes, were not detected after amino acid translation. Alignment of rDNA was done in BioEdit 7.2.1 (Hall, 1999) based on secondary structure information (Kjer, 1995) and previous studies (Pepato & Klimov, 2015, Pepato et al, 2018).

Best-fitting models of nucleotide substitutions were found in Partition Finder 1.0.1 (Lanfear *et al* 2012) using corrected Akaike Information Criterion (AICc). COI alignment was partitioned by codon positions and rDNA by paired (e.g., stems) and non-paired (e.g., loops) regions. Saturation of each partition was tested in DAMBE *6* (Xia et al., 2017); for CO1 the third codon position was excluded from analyses due to saturation (Iss > Iss.c: Iss = 1.239; Iss.cSym = 0.692 (P << 0.0001); Iss.cAsym = 0.364 (P << 0.0001)).

Molecular clock dating was performed in BEAST 2.3.2 (Bouckaert et al., 2014). Each analysis was run in five replicates and comprised at least $2 \times 10^8$ generations sampled every 10,000th generation. BEAST input file was generated in BEAUTi 2. Convergence of analyses was evaluated in Tracer 1.7 (Rambaut et al., 2018). Trees were visualized in FigTree 1.4.2 (http://tree.bio.ed.ac.uk/software/figtree/). Substitution models were set with the uncorrelated lognormal relaxed clock model (Drummond et al., 2006). The Yule speciation model showed a slightly better fit than the Birth and Death model using AICM, 289616.524 vs 289620.183 (calculated in Tracer). Because the former is less parameterized, it was preferred over the Birth and Death speciation model.

A lognormal prior distribution of ages was set to twelve paleontological calibration points associated with well-supported relationships, all treated as monophyletic clades (Table S2). Their offset corresponded to the minimal age and the mean calculated such that 95% of the distribution lies between the minimal ages and the soft maximum bounds, with a standard deviation of 1.00. Minimal ages correspond to the minimal estimated age of descent from a given node. It may be the absolute age of the fossiliferous strata or the minimum age of the Stage/Age to which the fossil was assigned according the International Chronostratigraphic Chart (http://www.stratigraphy.org/ICSchart/ChronostratChart2017-02.jpg, accessed June 06, 2017). Maximal soft bounds were based upon maximal age of the first occurrences of fossils attributed to a well-supported, more inclusive clade to which the node belongs. Preliminary, prior-only runs were performed to check the densities for all calibrated nodes and establish an exponential distribution to the root that reflects the fossil record. The exponential distribution on the root was established with the mean = 16.9 and offset = 405.0 Ma. Prior distributions obtained from the priors-only analysis are given in Fig. S1.

The divergence times are given as medians and 95% highest posterior densities (HPDs) as estimated in TreeAnnotator. In order to employ the distributions obtained in these analyses for our secondary calibration, stationary trees were also summarized as means, given that medians and means may be used to set lognormal distributions that provide better descriptions of the output than normal distributions (Morrison, 2008).

**Results and Discussion**

Our topology (Fig. S2A-B) is similar to previous analyses concerning Halacaridae ingroup phylogeny (Pepato et al., 2018). It differs, however, by grouping the genus *Anystis* as sister group of Halacaridae in a clade comprising all Anystina with the following topology ((*Andocaeculus*, *Erythracarus*) ((Halacaridae, *Anystis*), Parasitengona)). For the first time the traditional grouping of Anystina was recovered in molecular analyses, possibly due to improved sampling. The split between the Halacaridae+*Anystis* clade and Parasitengona was recovered 374.8, 319.3-432.9 Ma, a result that pushes back this divergence in time if compared to previous studies (321.5, 264.0-381.3 Ma Pepato et al., 2018; 320, 270-345 Ma, Dabert et al., 2016). In our analysis, the divergence between *Anystis* and Halacaridae occurred at 329.4, 265.0-393.2 Ma.

The internal topology of Halacaridae was identical to that reported in Pepato et al. (2018), except that it includes a representative of the freshwater genus *Porolohmannella* and excludes some terminals with large proportions of missing data. The first branching in the stem Halacaridae was the subfamily Lohmannellinae (*Scaptognathus*), occurring 269.1, 216.7-329.3 Ma, a result similar to that published previously (271.3, 221.7-324.2 Ma). Therefore, once the transition between land and sea occurred at some point after the split from *Anystis*, the values for the age of transition from the land to the sea are similar to those recovered in Pepato et al. (2018).

Despite not being the focus of this study, it is noteworthy that the freshwater genus *Porolohmannella* was recovered as sister group to *Limnohalacarus*. It narrowed the age of freshwater transition in this lineage: in previous analyses, the divergence between the freshwater lineages and their marine ancestors occurred 219.2, 165.9-274.6 Ma. In the new analyses, the values are very similar (224.3, 151.6-245.6 Ma), but now the divergence between *Porlohmannella* and *Limnohalacarus* was 136.8, 63.1-204.8 Ma, pushing back in time the minimal age for this freshwater lineage crown group.

The resemblance in gnathosomal morphology of Pezidae (a rare Australian lineage, not available for sequencing) and *Lohmannella* and *Porolohmannella*, was the main reason of placing the freshwater family Pezidae in Halacaroidea (Harvey, 1990). Because *Porolohmannella* does not form a monophyletic group with the remaining Lohmannellinae (as expected based on overall similarity) and considering that Pezidae females share a conspicuous apomorphic character (females attach their eggs to their hind legs), our new analysis suggests that Peziidae is a very derivative Halacaridae, and not a separate family. It would be very interesting to definitively solve this evolutionary question in the future.

Our focal taxon, the *Rhombognathus levigatoides* species complex was recovered as sister species diverging from each other 7.2, 1.69-16.28 Ma. It was recovered in a weakly supported clade with the undescribed species occurring in the South of São Paulo State (PP= 0.69), and hence was not employed as a secondary calibration in downstream analyses. These three terminals were grouped with an unnamed species from California (27.1, 14.37-43.12 Ma), and then with *Rhombognathus areolatus* (39.3, 21.7-60.39 Ma). *Agauopsis legionium* diverged from its sister species from California at 43.3 (17.6, 76.7), both diverging from *Agauopsis bilophus* at 74.5 (40.7, 114.8) Ma.

**Table S1**. Terminals, their taxonomic classification, and GenBank accession numbers for the nuclear ribosomal genes SSU (18S), LSU (28S) and mitochondrial locus COI. This dataset was used to infer our acariform mite time tree (Fig. 2S).

| Taxon<br>Order Solifugae | Family or Subfamily | SSU | LSU | COI |
|---|---|---|---|---|
| *Mummucia* | Mummucidae | KY922096 | KY921963 | KY922346 |
| *Gluvia dorsalis* | Daesiidae | AF007103 | KM100936 | KM100979 |
| *Eremobates* | Eremobatidae | AY859573 | AY859572 | ----- |
| Order Acariformes<br>Endeostigmata | | | | |
| *Nanorchestes* | Nanorchestidae | KY922106 | KY921973 | KY922356 |
| *Bimichaelia* | Alycidae | KY922112 | KY921980 | KY922358 |
| *Pachygnathus* | Alycidae | KY922115 | KY921983 | KY922360 |
| *Alycus aff roseus* | Alycidae | KY922116 | KY921984 | KY922361 |
| *Cunliffea* | Nematalycidae | KY922118 | KY921987 | KY922363 |
| *Gordialycus* | Nematalycidae | KY922131 | KY921999 | KY922376 |
| *Alicorhagia* | Alicorhagiidae | JQ000032 | JQ000339 | KY922381 |
| *Micropsammus* | Micropsammidae | KY922132 | KY922000 | KY922377 |
| *Terpnacarus gibbosus* | Terpnacaridae | KY922140 | KY922008 | KY922387 |
| Oehserchestidae | Oehserchestidae | KY922143 | KY922011 | KY922390 |
| *Hybalicus* | Lordalychidae | KY922103 | KY921970 | KY922353 |
| Oribatida<br>Palaeosomata | | | | |
| *Palaeacarus kamenskii* | Palaeacaridae | JQ000034 | JQ000341 | KY922451 |
| *Beklemishevia galeodula* | Ctenacaridae | KP325051 | KP325013 | ---- |
| Enarthronota | | | | |
| *Liochthonius* sp. *brevis*-complex | Brachychthoniidae | JQ000035 | JQ000342 | KY922452 |
| Brachychthoniidae sp | Brachychthoniidae | JQ000036 | JQ000343 | KY922453 |
| *Paralycus* | Pediculochelidae | KY922209 | KY922080 | KY922457 |
| *Cosmochthonius lanatus* | Cosmochthoniidae | JQ000040 | JQ000348 | ---- |
| *Hypochthonius luteus* | Hypochthoniidae | JQ000038 | JQ000345 | KY922455 |
| *Haplochthonius simplex* | Haplochthoniidae | KY922210 | JQ000347 | KY922458 |
| Mixonomata | | | | |
| *Euphthiracarus pulchrus* | Euphthiracaridae | JQ000041 | JQ000349 | KY922460 |
| *Meristacarus* sp. | Lohmaniidae | KP276478 | KP276398 | ---- |
| *Epilohmannia pallida* | Epilohmanniidae | KY922212 | KY922082 | KY922461 |
| Parhyposomata | | | | |
| *Gehypochthonius* | Gehypochthoniidae | JQ000042 | JQ000350 | KY922462 |
| Holosomata | | | | |
| *Nothrus borussicus* | Nothridae | KY922216 | KY922086 | KY922468 |
| *Camisia segnis* | Camisiidae | JQ000043 | JQ000351 | KY922463 |
| *Camisia* sp. | Camisiidae | MK014972 | MK014995 | MK015019 |
| *Masthermannia* sp | Nanhermanniidae | KY922217 | KY922087 | ---- |
| *Afronothrus* sp. | Trhypochthoniidae | JQ000045 | JQ000353 | KY922467 |
| *Trhypochthonius americanus* | Trhypochthoniidae | JQ000046 | JQ000354 | KY922469 |
| Brachypylina | | | | |
| *Trachyoribates ovulum* | Haplozetidae | HM070342 | KP276400 | ----- |
| *Gymnodamaeus bicostatus* | Gymnodamaeidae | MK014973 | MK014996 | MK015020 |
| *Chamobates* sp. | Chamobatidae | MK014974 | MK014997 | MK015021 |
| *Zetorchestes micronychus* | Zetorchestidae | MK014975 | MK014998 | MK015022 |
| *Pergalumna cf. nervosa* | Galumnidae | MK014976 | MK014999 | MK015023 |

| | | | | |
|---|---|---|---|---|
| *Neoribates aurantiacus* | Parakalummidae | MK014977 | MK015000 | MK015024 |
| *Cultroribula bicultrata* | Astegistidae | MK014978 | MK015001 | MK015025 |
| *Siculobata leontonycha* | Hemileiidae | MK014979 | MK015002 | MK015026 |
| *Aeroppia* sp. | Oppiidae | HM070344 | KP276401 | ----- |
| *Carabodes* sp. | Carabodidae | MK014980 | MK015003 | MK015027 |
| *Odontocepheus oblongus* | Carabodidae | KY922219 | KY922089 | KY922471 |
| *Cubabodes* aff *verrucatus* | Carabodidae | KY922220 | KY922090 | KY922472 |
| *Charassobates* aff *tuberosus* | Charassobatidae | KY922221 | KY922091 | KY922473 |
| *Anachipteria howardi* | Achipteriidae | JQ000048 | JQ000356 | ----- |
| *Pseudotocepheus amonstruosus* | Tetracondylidae | HM070341 | KP276403 | ----- |
| Astigmata "Acaridia" | | | | |
| *Naiadacarus arboricola* | Acaridae | JQ000114 | JQ000422 | KY922487 |
| *Canestrinia pentodontis* | Canestriniidae | JQ000084 | JQ000392 | ----- |
| *Dermacarus tamiasciuri* | Glycyphagidae | KP325070 | KP325032 | ----- |
| *Nanacarus* | Hemisarcoptidae | JQ000068 | JQ000376 | KY922477 |
| *Bonomoia* sp | Histiostomatidae | JQ000050 | JQ000358 | KY922474 |
| *Picidectes chapini* | Hypoderatidae | JQ000126 | JQ000434 | ----- |
| Psoroptidia | | | | |
| *Avenzoaria calidridis* | Avenzoariidae | KP325068 | KP325030 | KU203080 |
| *Dermatophagoides farinae* | Pyroglyphidae | JQ000247 | JQ000555 | GQ465336 |
| *Chirnyssoides amazonae* | Sarcoptidae | KP325067 | KP325029 | ----- |
| *Amerodectes aff atyeoi* | Proctophyllodidae | KP325069 | KP325031 | KU203226 |
| Prostigmata Eupodina | | | | |
| *Labidostomma* (*Atyeonella*) sp. | Labidostommatidae | KP276485 | KP276409 | ----- |
| Labidostomatidae gen. sp. | Labidostomatidae | KY922146 | KY922014 | KY922392 |
| *Scolotydeus uralensis* | Paratydeidae | MK014981 | MK015004 | MK015028 |
| *Tanytydeus* sp. | Paratydeidae | KY922147 | KY922015 | KY922393 |
| Ereynetidae gen. sp. | Ereynetidae | KY922152 | KY922020 | KY922398 |
| Tydeidae gen. sp. | Tydeidae | KY922148 | KY922016 | KY922394 |
| *Pseudolorrya* sp. | Tydeidae | MK014982 | MK015005 | MK015029 |
| *Tydeus* sp. | Tydeidae | MK014983 | MK015006 | MK015030 |
| *Cyta* sp. | Bdellidae | MK014985 | MK015008 | ---- |
| *Bdella longicornis* | Bdellidae | KY922154 | KY922023 | KY922401 |
| *Parabonzia* sp. | Cunaxidae | KY922155 | KY922024 | KY922402 |
| *Benoinyssus aff serratus* | Eupodidae | KY922150 | KY922018 | KY922396 |
| *Linopodes* sp. | Eupodidae | KY922153 | KY922022 | KY922400 |
| Eupodidae gen. sp. | Eupodidae | MK014984 | MK015007 | MK015031 |
| Rhagidiidae gen. sp. | Rhagidiidae | KY922151 | KY922019 | ---- |
| *Penthalodes sp.* | Penthalodidae | KP276486 | KP276410 | ---- |
| *Stereotydeus* sp | Penthalodidae | KP276487 | KP276411 | ---- |
| Eleutherengona | | | | |
| *Eustigmaeus* sp. | Stigmaeidae | KY922175 | KY922046 | KY922420 |
| *Homocaligus* sp. | Homocaligidae | KY922176 | KY922047 | KY922421 |

| | | | | |
|---|---|---|---|---|
| *Caenolestomyobia faini* | Myobiidae | KY922177 | KY922048 | KY922422 |
| *Radfordia elegantula* | Myobiidae | KY922178 | KY922049 | KY922423 |
| *Harpyrhynchoides zumpti* | Harpirhynchidae | KY922181 | KY922052 | KY922426 |
| *Harpypalpus holopus* | Harpirhynchidae | KY922185 | KY922056 | KY922429 |
| *Psorergates* sp. | Psorergatidae | KY922186 | KY922057 | KY922430 |
| *Demodex folliculorum* | Demodecidae | KY922187 | KY922058 | KY922431 |
| *Barbutia* sp. | Barbutiidae | KY922188 | KY922059 | KY922432 |
| *Cyclurobia* sp. | Pterygosomatidae | KY922190 | KY922061 | KY922434 |
| Caligonellidae gen. sp. | Caligonellidae | KY922191 | KY922062 | KY922435 |
| *Raphignathus* sp. | Raphignathidae | KY922192 | KY922063 | KY922436 |
| *Favognathus* sp. | Cryptognathidae | KY922193 | KY922064 | KY922437 |
| Linotetranidae gen. sp. | Linotetranidae | KY922194 | KY922065 | KY922438 |
| *Eotetranychus* sp. | Tetranychidae | KY922195 | KY922066 | KY922439 |
| *Tetranychus urticae* | Tetranychidae | KY922197 | KY922068 | KY922441 |
| *Tenuipalpus* sp. | Tenuipalpidae | KY922198 | KY922069 | KY922442 |
| *Brevipalpus* sp. | Tenuipalpidae | KY922199 | KY922070 | KY922443 |
| *Neophyllobius* sp. | Camerobiidae | KY922200 | KY922071 | KY922444 |
| *Bakericheyla chanayi* | Cheyletidae | KY922201 | KY922072 | KY922445 |
| *Chelacheles* sp. | Cheyletidae | KY922202 | KY922073 | KY922446 |
| Cheyletidae gen. sp. | Cheyletidae | KY922203 | KY922074 | KY922447 |
| *Oudemansicheyla* sp. | Cheyletidae | HM070362 | KP276422 | ----- |
| *Charadriphilus gallinago* | Syringophilidae | KY922205 | KY922076 | KY922448 |
| Anystina | | | | |
| *Andocaeculus* sp. | Caeculidae | KP276488 | KP276412 | ----- |
| *Erythracarus* sp. | Anystidae | KP276489 | KP276413 | ----- |
| *Anystis* sp. | Anystidae | KY922145 | KP325014 | KY922391 |
| Adamystidae gen. sp. | Adamystidae | KY922156 | KY922026 | KY922403 |
| *Eutrombicula splendens* | Trombiculidae | KY922159 | KY922031 | MK015032 |
| *Acomatacarus arizonensis* | Leeuwenhoekiidae | KY922157 | KY922029 | KY922406 |
| *Leptus* sp. | Erythraeidae | KP276490 | KP276414 | ----- |
| *Caeculisoma* sp. | Erythraeidae | KP276491 | KP276415 | ----- |
| *Balaustium* sp. | Erythraeidae | KY922158 | KY922030 | KY922407 |
| *Lasioerythraeus* sp. | Erythraeidae | KM100884 | KM100950 | KM100991 |
| *Smaris sp.* | Smarididae | KM100885 | KM100951 | KM100990 |
| *Calyptostoma* sp. | Calyptostomatidae | KM100878 | KM100948 | KM100992 |
| *Calyptostoma velutinus* | Calyptostomatidae | KM100880 | KM100949 | KM100993 |
| *Diplothrombium* sp. | Johnstonianidae | KM100930 | KM100940 | KM100986 |
| *Allothrombidium* sp. | Trombidiidae | KP276493 | KP276417 | ---- |
| *Dactylothrombium pulcherrimum* | Microtrombidiidae | GQ864281 | KM100939 | KM100985 |
| *Valgoperuvia paradoxa* | Microtrombidiidae | KM100934 | KM100943 | KM100988 |
| *Stygothrombium* sp. | Stygothrombidiidae | KM100927 | KM100938 | KM100995 |
| *Arrenurus (A.)* sp. | Arrenuridae | KM100875 | KM100944 | KM101003 |
| *Horreolanus orphanus* | Bogatiidae | AY620907 | KM100946 | KM101004 |
| *Mideopsis roztoczensis* | Mideopsidae | JN018219 | JN018316 | JN018102 |
| *Eylais* sp | Eylaidae | KM100887 | KM100955 | AY393896 |
| *Limnochares americana* | Limnocharidae | KM100888 | KM100956 | KM100998 |
| *Hydrachna conjecta* | Hydrachnidae | JN018220 | JN018317 | JN018103 |
| *Hydrovolzia placophora* | Hydrovolziidae | KM100889 | KM100957 | KM100996 |

| | | | | |
|---|---|---|---|---|
| *Hydryphantes waynensis* | Hydryphantidae | KM100893 | KM100959 | KM101012 |
| *Wandesia* sp. | Hydryphantidae | KM100897 | KM100960 | KM101010 |
| *Feltria* sp. | Feltriidae | KM100905 | KM100965 | KM101021 |
| *Coaustraliobates cf. cortipes* | Hygrobatidae | KM100904 | KM100964 | KM101024 |
| *Limnesia (Limnesiella) marshallae* | Limnesiidae | KM100910 | KM100969 | KM101029 |
| *Australotiphys barmutai* | Pionidae | KM100903 | KM100963 | KM101017 |
| *Litarachna communis* | Pontarachnidae | KM100911 | KM100970 | KM101025 |
| *Unionicola crassipes* | Unionicolidae | KM100918 | KM100973 | KM101023 |
| *Frontipoda* sp. | Oxidae | KM100919 | KM100975 | KM101000 |
| *Sperchonopsis phreaticus* | Sperchontidae | KM100922 | KM100978 | KM100999 |
| *Teutonia cometes* | Teutoniidae | JN018224 | JN018321 | JN018107 |
| *Torrenticola amplexa* | Torrenticolidae | JN018226 | JN018323 | JN018191 |
| *Halacaroides antoniazae.* | Halacarinae | MG751443 | MG751417 | MG696228 |
| *Acarothrix* sp. | Copidognathinae | KP276481 | KP276405 | MG696250 |
| *Copidognathus* sp1 | Copidognathinae | MG751444 | MG751418 | MG696229 |
| *Copidognathus* sp2 | Copidognathinae | MG751445 | MG751419 | MG696230 |
| *Copidognathus* sp3 | Copidognathinae | MG751446 | MG751420 | MG696231 |
| *Copidognathus* sp.5 | Copidognathinae | MK014986 | MK015009 | ---- |
| *Agauopsis legionium* | Halacarinae | MG751447 | MG751421 | MG696232 |
| *Agauopsis bilophus* | Halacarinae | MG751448 | MG751422 | MG696233 |
| Agauopsis sp. | Halacarinae | MK014987 | MK015010 | MK015033 |
| *Halacarellus* sp. | Halacarinae | MG751449 | MG751423 | MG696234 |
| *Halacaropsis* cf *hirsuta* | Halacarinae | MG751450 | MG751424 | MG696235 |
| *Halacarus omului* | Halacarinae | MG751451 | MG751425 | MG696236 |
| *Thalassarachna sp.* | Halacarinae | MG751452 | MG751426 | MG696237 |
| *Agaue* sp. | Halixodinae | MG751453 | MG751427 | MG696238 |
| *Bradyagaue sp.* | Halixodinae | MG751454 | MG751428 | MG696239 |
| *Limnohalacarus cultellatus* | Limnohalacarinae | MG751455 | MG751429 | ----- |
| *Limnohalacarus mamillatus*. | Limnohalacarinae | KP276482 | KP276406 | ----- |
| *Scaptognathus* sp1 | Lohmannellinae | MG751457 | MG751431 | MG696240 |
| *Scaptognathus* sp2 | Lohmannellinae | MG751458 | MG751432 | MG696241 |
| *Porolohmannella violacea* | Porolohmannellinae | MK014988 | MK015011 | MK015034 |
| *Isobactrus setosus* | Rhombognathinae | MG751460 | MG751434 | ----- |
| *Isobactrus uniscutatus* | Rhombognathinae | MG751461 | MG751435 | MG696243 |
| *Isobactrus* sp | Rhombognathinae | MK014989 | MK015012 | MK015035 |
| *Metarhombognathus armatus* | Rhombognathinae | KP276483 | KP276407 | MG696251 |
| *Rhombognathus* sp.1 | Rhombognathinae | MK014990 | MK015013 | MK015036 |
| *Rhombognathus areolatus* | Rhombognathinae | MG751463 | MG751437 | MG696244 |
| *Rhombognathus levigatoides* (SE) | Rhombognathinae | MG751464 | MG751438 | MG696245 |
| *Rhombognathus levigatoides* (NE) | Rhombognathinae | MK014991 | MK015014 | MK015037 |
| *Rhombognathus* sp. 2 | Rhombognathinae | MK014992 | MK015015 | MK015038 |

| | | | | |
|---|---|---|---|---|
| *Rhombognathus* sp. 3 | Rhombognathinae | MK014993 | MK015016 | MK015039 |
| *Rhombognathus* sp. 4 | Rhombognathinae | MG751465 | MG751439 | MG696246 |
| *Acaromantis vespucioi* | Simognathinae | MG751466 | MG751440 | MG696247 |
| *Simognathus* sp1 | Simognathinae | MG751467 | MG751441 | MG696248 |
| *Simognathus* sp2 | Simognathinae | MG751468 | MG751442 | MG696249 |
| *Simognathus* sp.3 | Simognathinae | MK014994 | MK015017 | MK015040 |

**Table S2** Fossil calibration points used in BEAST molecular clock phylogenetic analyses. Lognormal prior distribution of ages was set with an offset corresponding to the minimal age and the mean calculated such that 95% of the distribution lies between the minimal ages and the maximum age, with a standard deviation of 1.00. The only exception was the root: its prior distribution was set as an exponential distribution by running preliminary priors-only analyses (Fig. S1) to check whether the prior fossil calibration distributions are affected by priors' interactions. Lineages represent crown groups, unless otherwise indicated.

| **Divergence** | **Min. age** | **Max. age** | **Fossil** | **Reference** |
|---|---|---|---|---|
| **Root** | 405.1 | 514.0 | Min.: *Protacarus crani*<br>Max.: *Wisangocaris barbarahardyae* | Wolfe et al., 2016 |
| **Solifugae** | 112.6 | 509.0 | Min.: *Cratosolpuga wunderlichi*<br>Max.: *Wisangocaris barbarahardyae* | Wolfe et al., 2016 |
| **Sarcoptiformes** | 405.1 | 509.0 | Min.: *Protospeleorchestes pseudoprotacarus*;<br>Max.: *Wisangocaris barbarahardyae* | Dubinin, 1962; Wolfe et al., 2016 |
| **Erythraeoidea** | 112.6 | 410.2 | Min.: *Pararainbowia martilli*<br>Max.: *Protacarus crani* | Dunlop, 2007 |
| **Pterygosomatidae** | 100.5 | 410.2 | Min.: *Pimeliaphilus* sp.<br>Max.: *Wisangocaris barbarahardyae* | Sidorchuk & Khaustov, 2018 |
| **Cheyletidae** | 98.2 | 410.2 | Min.: *Cheyletus burmiticus*<br>Max: *Protacarus crani* | Cockerell, 1917 |
| **stem group Enarthronota** | 388.1 | 509.0 | Min.: *Protochthonius gilboa, Devonacarus sellnicki*<br>Máx.: *Wisangocaris barbarahardyae* | Norton et al., 1988 |

| | | | | |
|---|---|---|---|---|
| **Lohmanniidae-Hypochthoniidae** | 326.4 | 410.2 | Min.: *Palaeohypochthonius jerami*<br>Max.: *Protacarus crani* | Subías & Arillo, 2002 |
| **Brachypilina** | 189.8 | 388.1 | Min.: *Hydrozetes* sp.<br>Max.: *Protochthonius gilboa*, *Devonacarus sellnicki* | Sivhed & Wallwork, 1978 |
| **Trhypochthoniidae** | 100.5 | 388.1 | Min.:*Trhypochthonius lopezvallei*<br>Max.: *Protochthonius gilboa*, *Devonacarus sellnicki* | Arillo et al., 2012 |
| **Astegistidae** | 145.0 | 388.1 | Min.: *Cultroribula jurassica*<br>Max.: *Protochthonius gilboa*, *Devonacarus sellnicki* | Krivolutsky & Krasilov, 1977 |
| **Achipteriidae** | 145.0 | 388.1 | Min.: *Achipteria obscura*<br>Max.: *Protochthonius gilboa*, *Devonacarus sellnicki* | Krivolutsky & Krasilov, 1977 |

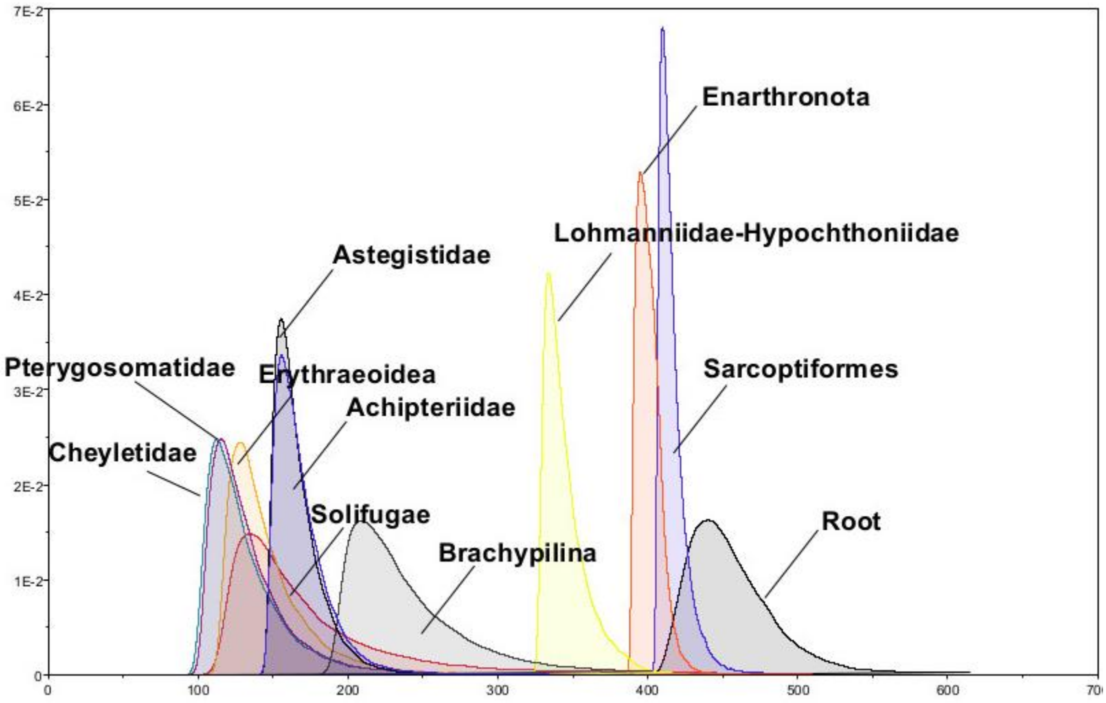


**Figure S1**. Multiplicative priors on node ages estimated by Tracer. See Table 2 for a list of calibration points and min/max ages. Age distributions represent crown groups, except for Enarthronota (stem group). [label y and x axes]

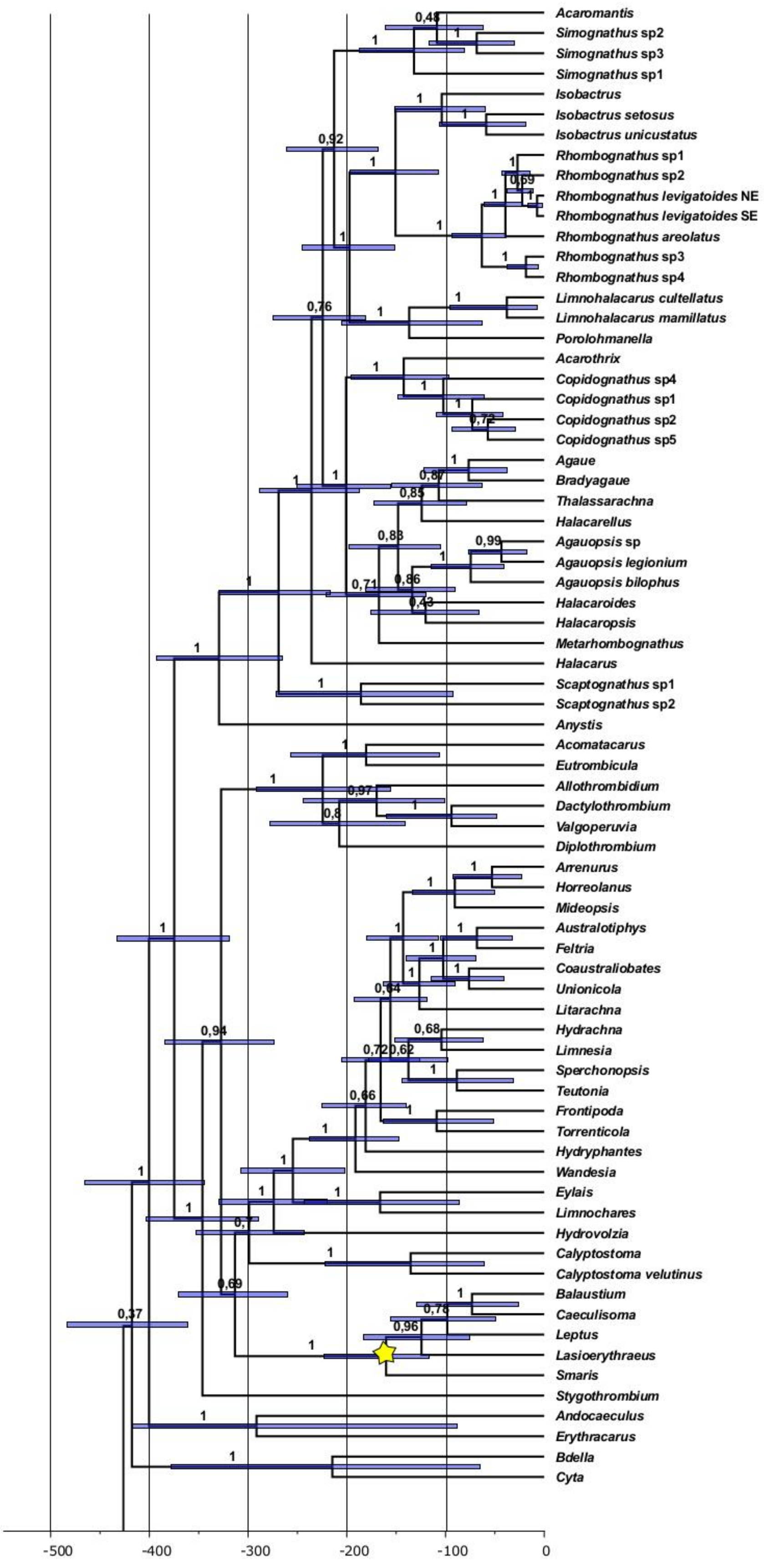


Figure S2A. Time calibrated tree for Acariformes with fossil calibration points indicated by stars. Values above branches are posterior probabilities; bars represent 95% HPD interval.

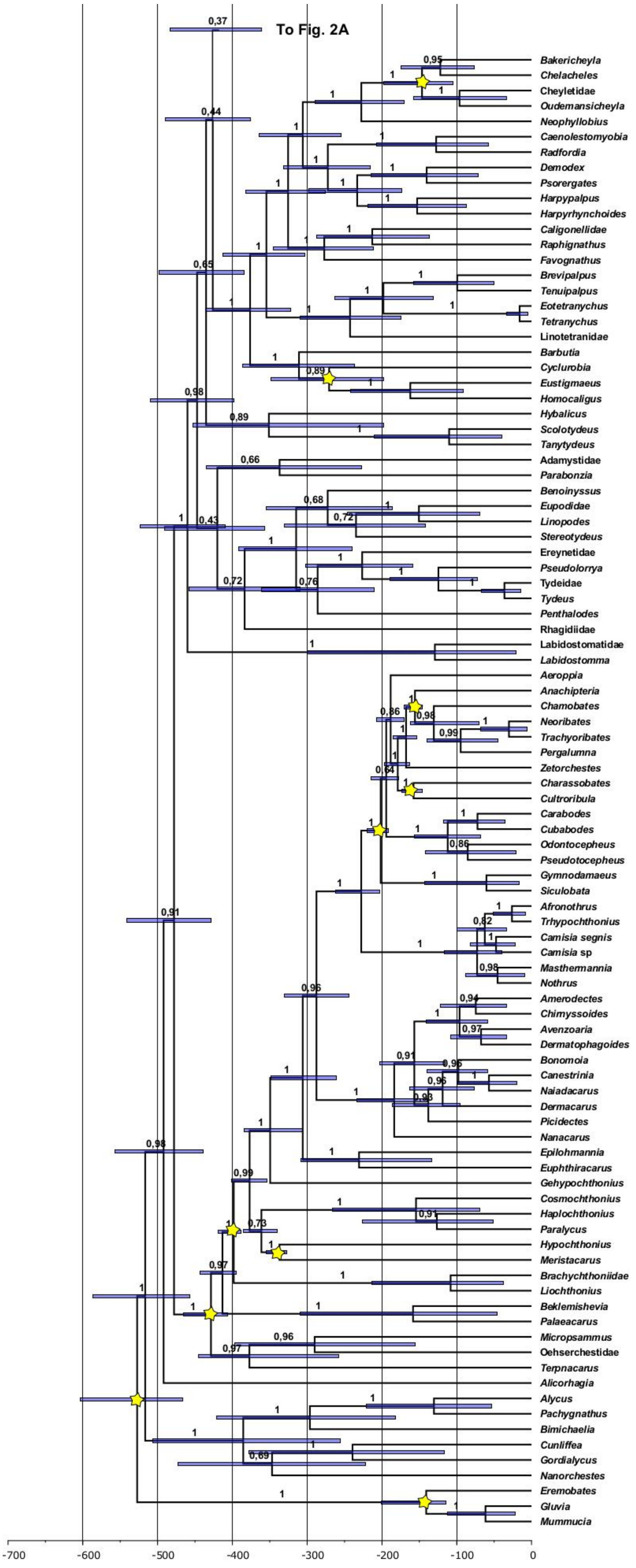


Figure S2B (Continues from Fig. S2A). Time calibrated tree for Acariformes mites with fossil calibration points indicated by stars. Values above branches are posterior probabilities; bars are the 95% HPD interval.
