## Appendix 4 for "Evaluating the boundaries of marine biogeographic regions of the Southwestern Atlantic using halacarid mites (Halacaridae), meiobenthic organisms with a low dispersal potential"

**Appendix 5: Niche comparison and modeling**

In this appendix, we present the results from some analyses accessory to those presented in the main text and provide the R scripts employed to perform them. It is largely based on R package manuals and vignettes, but we consider that presenting them here may help future researchers interested in using these packages.

Script 1. R script for checking layers for correlation.

```
#For installing the ENMTools package (skip if already installed)
install.packages("devtools")
library(devtools)
install_github("danlwarren/ENMTools")
#load libraries
library(ENMTools)
#Create a stack from the ascii files. Adjust the path to the folder according to your computer
env.files <- list.files(path = "C:/Users/Dell/Documents/layers", full.names = TRUE)
env <- stack(env.files)
crs(env) <- " +proj=longlat +datum=WGS84 +ellps=WGS84 +towgs84=0,0,0"
#It is optional, you may want plot the layers to check them individually
plot(env)
#Calculate the Pearson Correlation Matrix
cor_matrix<-raster.cor.matrix(env)
#Plot a heat map with the results
cor_plots<-raster.cor.plot(env)
#write a file with a table including the values
write.table(cor_matrix, file = "table_correlation.csv")
```

Table 1. Layers considered for Enviromental Niche Modeling analyses and highly correlated (Pearson correlation coefficient module larger than 0.75 ) variables excluded from further consideration.

| Included Layers (>0.75) | Definition and hightly correlated Layers |
|---|---|
| Bathymetry (MARSPEC) | Depth of the seafloor (m). |
| Bio2 (ECOCLIMATE) | Mean diurnal range (°C) (mean of monthly (max temp min temp)). |
| Bio3 (ECOCLIMATE) | Isothermality (%) (100*Bio2/Bio7). |
| Bio12 (ECOCLIMATE) | Annual precipitation (mm/m$^2$). Bio13 (ECOCLIMATE): Precipitation of wettest month (mm/m$^2$), Bio16 (ECOCLIMATE): Precipitation of wettest quarter (mm/m$^2$). |
| Bio18 (ECOCLIMATE) | Precipitation of warmest quarter (mm/m$^2$) |
| Bio19 (ECOCLIMATE) | Precipitation of coldest quarter (mm/m$^2$) |
| Biogeo1 (MARSPEC) | East/West Aspect (radians) |
| Biogeo2 (MARSPEC) | North/South Aspect (radians) |
| Biogeo3 (MARSPEC) | Plan Curvature |
| Biogeo4 (MARSPEC) | Profile Curvature; Biogeo7 (MARSPEC) Concavity (degrees); Biogeo6 (MARSPEC): Bathymetric Slope (degrees); |
| Biogeo9 (MARSPEC) | Sea Surface Salinity (SSS) of the freshest month (psu). Biogeo8 (MARSPEC): Mean Annual SSS (psu); Biogeo11, Annual range in SSS (psu) Salinity maxn (Bio-Oracle) (PSS); Salinity mean (Bio-Oracle) (PSS); Chlorophyll A min. (Bio-Oracle); Chlorophyll A min(mg/m$^3$). Phytoplankton min (mmol/m$^3$); Primary productivity mean (g/m$^3$day); Primary productivity min (g/m$^3$day); Silicate max. (Bio-Oracle): Silicate max. concentration (mmol/m$^3$). Biogeo11 (MARSPEC): |

| | |
|---|---|
| | Annual range in SSS (psu); Biogeo12 (MARSPEC): Annual variance in SSS (psu); Dissolved Iron max. (mmol/$m^3$); Dissolved Iron mean (mmol/$m^3$); Dissolved Iron min. (mmol/$m^3$); Nitrate concentration mean (mmol/$m^3$); Salinity mean (Bio-Oracle) (PSS); Salinity min (Bio-Oracle) (PSS); Silicate min concentration (Bio-Oracle) (mmol/$m^3$); Silicate mean concentration (Bio-Oracle) (mmol/$m^3$); pH (Bio-Oracle) Sea water pH. |
| Biogeo10 (MARSPEC) | SSS of the saltiest month (psu) |
| Calcite concentration (Bio-Oracle) | Mean Calcite concentration (mol/$m^3$) |
| Chlorophyll A mean (Bio-Oracle) | Chlorophyll A mean (mg/$m^3$). Chlorophyll A max (Bio-Oracle), Phytoplankton max (mmol/$m^3$); Phytoplankton mean (mmol/$m^3$); Primary productivity max (g/$m^3$day). |
| Cloud cover mean (Bio-Oracle) | Cloud cover mean (%). Cloud cover max (Bio-Oracle): Cloud cover max (%). |
| Cloud cover min. | Cloud cover min (%). |
| Current velocity max. (Bio-Oracle) | Current velocity max (m/s). |
| Current velocity mean (Bio-Oracle) | Current velocity mean (m/s). Current velocity min (m/s) (Bio-Oracle) |
| Diffuse attenuation max. (Bio-Oracle) | Diffuse attenuation max. ($m^{-1}$). Diffuse attenuation min. ($m^{-1}$). |
| Light at the bottom max. (Bio-Oracle) | Light at the bottom max. (Einstein/$m^2$day). |
| Light at the bottom min. (Bio-Oracle) | Light at the bottom min. (Einstein/$m^2$day). |
| Nitrate min. (Bio-Oracle) | Nitrate concentration min. (mmol/$m^3$). Silicate concentration min. (mmol/$m^3$); Salinity max. (Bio-Oracle ) (PSS). Salinity mean (Bio-Oracle (PSS). |
| Photosynthetically Active radiation max (Bio-Oracle) | Photosynthetically Active radiation max (Einstein/$m^2$/day) |
| Sea Surface Temperature mean (Bio-Oracle) | Sea Surface Temperature mean (°C). Bio1 (ECOCLIMATE): Annual mean temperature (°C); Bio10 (ECOCLIMATE): Mean temperature of warmest quarter (°C); Bio11 (ECOCLIMATE): Mean temperature of coldest quarter (°C); Bio14 (ECOCLIMATE): Precipitation of driest month (mm/m2); Bio15 (ECOCLIMATE): Precipitation seasonality - % (coefficient of variation); Bio 4 (ECOCLIMATE): Temperature seasonality (%) (standard deviation *100); Bio9 (ECOCLIMATE): Mean temperature of driest quarter (°C); Bio17 (ECOCLIMATE): Precipitation of driest quarter (mm/$m^2$); Biogeo13 (MARSPEC): Mean Annual Sea Surface Temperature (SST, °C); Biogeo14 (MARSPEC): SST of the coldest month (°C); Biogeo15 (MARSPEC): SST of the warmest month (°C); biogeo16 (MARSPEC): Annual range in SST (°C); Biogeo17 (MARSPEC): Annual variance in Sea Surface Temperature (SST, °C); Dissolved molecular oxygen max. (mmol/m3), Photosynthetically Active radiation mean (Einstein/$m^2$day); Sea Surface Temperature max (Bio-Oracle) (°C); Sea Surface Temperature min (Bio-Oracle) (°C). Bio6 (ECOCLIMATE): Min temperature of coldest month (°C); Bio5 (ECOCLIMATE): Max temperature of warmest month (°C); Bio7 (ECOCLIMATE): Temperature annual range (°C) (Bio5-Bio6) Bio8 (ECOCLIMATE): Mean temperature of wettest quarter (°C); Phosphate mean (Bio-Oracle): Phosphate mean concentration (μmol/l). Phosphate max. concentration (μmol/l); Phosphate min. concentration (μmol/l); Dissolved molecular oxygen min. (Bio-Oracle): Dissolved molecular oxygen min. (mmol/m3). |

Script 2. R script for pre-modelling niche comparison

```
#load libraries
library (ecospat)
library (ENMTools)
library (raster)
#Occurrence data
# Load the occurrence records. In our case, both sibling species of Rhombognathus
LonLatDataNE <- read.csv("Rhombognathus_NE.csv")[,2:3]
LonLatDataSE <- read.csv("Rhombognathus_SE.csv")[,2:3]
#Then load the environmental variables into R with the help of the stack function of the 'raster' package.
#You can not just copy the following line but have to adjust the filepath to your own.
files <- list.files("C:/Users/Dell/Documents/layers.75",pattern='asc',full.names=TRUE)
Grids <- raster::stack(files)
#Create a background for each species to emulate a known distribution
env.files <- list.files(path = "C:/Users/Dell/Documents/layers.75", full.names = TRUE)
env <- stack(env.files)
env <- setMinMax(env)
crs(env) <- " +proj=longlat +datum=WGS84 +ellps=WGS84 +towgs84=0,0,0"
NE_SP <- enmtools.species()
NE_SP$species.name <- "NE"
NE_SP$presence.points <- LonLatDataNE
NE_SP$range <- background.raster.buffer(NE_SP$presence.points, 20000, mask = env)
NE_SP$background.points <- background.points.buffer(points = NE_SP$presence.points, radius = 20000,
n = 500, mask = env[[1]])
#take a look on the results
plot (NE_SP)
SE_SP <- enmtools.species()
SE_SP$species.name <- "SE"
SE_SP$presence.points <- LonLatDataSE
SE_SP$range <- background.raster.buffer(SE_SP$presence.points, 20000, mask = env)
SE_SP$background.points <- background.points.buffer(points = SE_SP$presence.points, radius = 20000, n
= 500, mask = env[[1]])
#take a look on the results
plot (SE_SP)
#Extract variables from each species range
VariablesAtNE_SPbg <- raster::extract(Grids,NE_SP$background.points)
VariablesAtSE_SPbg <- raster::extract(Grids,SE_SP$background.points)
#Values for background points for each species
OutputNE_SPbg <- as.data.frame(cbind("species", NE_SP$background.points, VariablesAtNE_SPbg))
colnames(OutputNE_SPbg) <- c("species","x","y", colnames(VariablesAtNE_SPbg))
OutputSE_SPbg <- as.data.frame(cbind("species", SE_SP$background.points, VariablesAtSE_SPbg))
colnames(OutputSE_SPbg) <- c("species","x","y", colnames(VariablesAtSE_SPbg))
# Extracting the variables for all occurrencelocations
VariablesAtOccurrencelocationsNE <- raster::extract(Grids,LonLatDataNE)
VariablesAtOccurrencelocationsSE <- raster::extract(Grids,LonLatDataSE)
# Combining the extracted values with the longitude and latitude values
NE_POINTS <- as.data.frame(cbind("Rhombognathus_NE", LonLatDataNE,
VariablesAtOccurrencelocationsNE))
SE_POINTS <- as.data.frame(cbind("Rhombognathus_SE", LonLatDataSE,
VariablesAtOccurrencelocationsSE))
colnames(NE_POINTS) <- c("species","x","y", colnames(VariablesAtOccurrencelocationsNE))
colnames(SE_POINTS) <- c("species","x","y", colnames(VariablesAtOccurrencelocationsSE))
#Complete dataset
NE<-rbind(NE_POINTS,OutputNE_SPbg)
NE <-NE[complete.cases(NE), ]
SE<-rbind(SE_POINTS,OutputSE_SPbg)
```

```
SE <-SE[complete.cases(SE), ]
#PCA.env score NE, SE and niche comparison
pca.env <- dudi.pca(rbind(NE,SE)[,4:ncol(NE)],scannf=F,nf=2)
#plot a correlation circle with PCA results
ecospat.plot.contrib(contrib=pca.env$co, eigen=pca.env$eig)
#Save variables correlations to principal components
write.table(pca.env$co, file = "corr_pca.csv")
# PCA scores for the whole study area
scores.globclim <- pca.env$li
# PCA scores for the species NE distribution
scores.sp.NE <- suprow(pca.env,NE[which(NE[,1]=="Rhombognathus_NE"),4:ncol(NE)])$li
# PCA scores for the species SE distribution
scores.sp.SE <- suprow(pca.env,SE[which(SE[,1]=="Rhombognathus_SE"),4:ncol(SE)])$li
# PCA scores for the whole NE study area
scores.clim.NE <- suprow(pca.env,NE[,4:ncol(NE)])$li
# PCA scores for the whole SE study area
scores.clim.SE <- suprow(pca.env,SE[,4:ncol(SE)])$li
# gridding the NE niche
grid.clim.NE <- ecospat.grid.clim.dyn(glob=scores.globclim,
glob1=scores.clim.NE,
sp=scores.sp.NE, R=100,
th.sp=0)
# gridding the SE niche
grid.clim.SE <- ecospat.grid.clim.dyn(glob=scores.globclim,
glob1=scores.clim.SE,
sp=scores.sp.SE, R=100,
th.sp=0)
# Compute Schoener's D, index of niche overlap
D.overlap <- ecospat.niche.overlap (grid.clim.NE, grid.clim.SE, cor=T)$D
D.overlap
#Delimiting niche categories and quantifying niche dynamics in analogue climates
#with ecospat.niche.dyn.index()
niche.dyn <- ecospat.niche.dyn.index (grid.clim.NE, grid.clim.SE, intersection = 0.1)
ecospat.plot.niche.dyn(grid.clim.NE, grid.clim.SE, quant=0.25, interest=2,
title= "Niche Overlap", name.axis1="PC1",
name.axis2="PC2")
ecospat.shift.centroids(scores.sp.NE, scores.sp.SE, scores.clim.NE, scores.clim.SE)
#Perform the Niche Equivalency Test according to Warren et al. (2008)
#Niche equivalency test H1: Is the overlap between the NE and SE niches higher than two random niches?
eq.testNESE <- ecospat.niche.equivalency.test(grid.clim.NE, grid.clim.SE,
rep=100, alternative = "greater")
ecospat.plot.overlap.test(eq.testNESE, "D", "Equivalency")
```

Table 2. Correlation of each variable relative to the two first Principal Components of A. legionium X *R. levigatoides* Southeastern Clade (Component 1 = 21.80 %, Component 2 = 18.65 % of variability), A. legionium X *R. levigatoides* Northeastern Clade (Component 1 = 19.83 %, Component 2 = 19.12 % of variability), and Southeastern and Northeastern of *R. levigatoides* complex (Component 1 = 23.92 %, Component 2 = 17.53 % of variability).

| | *A. legionium X R. levigatoides* SE | | *A. legionium X R. levigatoides* NE | | *R. levigatoides* NEXSE | |
|---|---|---|---|---|---|---|
| | Comp1 | Comp2 | Comp1 | Comp2 | Comp1 | Comp2 |
| Bathymetry | -0,55 | -0,13 | -0,51 | -0,43 | 0,02 | -0,71 |
| Mean diurnal range | -0,21 | -0,24 | -0,42 | -0,25 | 0,03 | -0,40 |
| Isothermality | -0,46 | -0,76 | -0,88 | -0,03 | -0,57 | -0,54 |
| Annual precipitation | -0,85 | 0,12 | -0,54 | -0,59 | 0,32 | -0,69 |
| Precipitation of warmest quarter | -0,49 | 0,31 | -0,22 | -0,06 | -0,01 | -0,47 |
| Precipitation of coldest quarter | -0,76 | 0,19 | -0,22 | -0,19 | 0,04 | -0,24 |
| East/West Aspect | 0,05 | 0,20 | 0,11 | 0,30 | -0,13 | -0,10 |
| North/South Aspect | -0,63 | -0,29 | -0,66 | -0,17 | -0,20 | -0,52 |
| Plan Curvature | -0,09 | 0,13 | -0,19 | -0,27 | 0,12 | -0,32 |
| Profile Curvature | -0,08 | 0,09 | -0,28 | -0,38 | 0,11 | -0,40 |
| Sea Surface Salinity of freshest month | 0,71 | -0,42 | 0,04 | 0,69 | -0,85 | 0,25 |
| SSS of the saltiest month | 0,09 | -0,73 | -0,34 | 0,45 | -0,79 | 0,39 |
| Mean Calcite concentration | -0,59 | -0,36 | -0,49 | -0,48 | 0,04 | -0,54 |
| Chlorophyll A mean | -0,02 | 0,65 | 0,45 | -0,69 | 0,86 | 0,03 |
| Cloud cover mean | -0,65 | -0,07 | -0,22 | 0,11 | -0,35 | -0,31 |
| Cloud cover min. | -0,43 | 0,06 | 0,00 | 0,36 | -0,33 | -0,12 |
| Current velocity max. | -0,23 | -0,38 | -0,48 | 0,62 | -0,59 | -0,40 |
| Current velocity mean | 0,14 | -0,70 | -0,55 | 0,31 | -0,62 | -0,15 |
| Diffuse attenuation max. | -0,80 | 0,08 | -0,33 | -0,72 | 0,48 | -0,56 |
| Light at the bottom max. | -0,35 | -0,07 | -0,33 | -0,24 | 0,03 | -0,54 |
| Light at the bottom min. | -0,31 | -0,54 | -0,71 | 0,04 | -0,46 | -0,67 |
| Nitrate min. | -0,40 | 0,56 | 0,28 | -0,47 | 0,60 | -0,20 |
| Photosynthetically Active radiation max | 0,36 | -0,48 | -0,19 | 0,77 | -0,73 | 0,15 |
| Sea Surface Temperature mean | -0,19 | -0,89 | -0,78 | 0,46 | -0,91 | -0,16 |

Script 3. R script for setting the feature numbers and regularization

In this case, note that we performed two classes of analyses, with and without shaping the background according to the sampling bias.

```
#load libraries
library (ENMeval)
library(ENMTools)
#read enviromental layers
env.files.NE <- list.files(path = "C:/Users/Dell/Documents/layers.NE", full.names = TRUE)
env.NE <- stack(env.files.NE)
env.NE <- setMinMax(env.NE)
crs(env.NE) <- " +proj=longlat +datum=WGS84 +ellps=WGS84 +towgs84=0,0,0"
#If you want take a look on them
plot(env.NE)
```

```
#read occurrences
Rhus_NE_occ <- read.csv("Rhombognathus_NE.csv")[,2:3]
#evaluate NE, no sampling bias (You may look at manual to change according your #application)
enmeval_Rhus_NE <- ENMevaluate(Rhus_NE_occ, env.NE, method="jackknife", n.bg=500, RMvalues =
seq(0.5, 3, 0.5), fc = c("L", "LQ"), algorithm='maxent.jar')
#Saving results
par(mfrow=c(2,2))
eval.plot(enmeval_Rhus_NE@results, "AICc")
dev.copy(pdf,'AICc_enmeval_Rhus_NE.pdf')
dev.off()
eval.plot(enmeval_Rhus_NE@results, "avg.test.AUC", variance="var.test.AUC")
dev.copy(pdf,'avgtestAUC_enmeval_Rhus_NE.pdf')
dev.off()
aic.mod <- enmeval_Rhus_NE@models[[which(enmeval_Rhus_NE@results$delta.AICc==0)]]
enmeval_Rhus_NE_varimpbest<-var.importance(aic.mod)
write.table(enmeval_Rhus_NE_varimpbest, file = "enmeval_Rhus_NE_varimpbest.csv")
enmeval_Rhus_NE_varimp<-lapply(enmeval_Rhus_NE@models, var.importance)
write.table(enmeval_Rhus_NE_varimp, file = "enmeval_Rhus_NE_varimp.csv")
plot(enmeval_Rhus_NE@predictions[[which (enmeval_Rhus_NE@results$delta.AICc == 0) ]])
dev.copy(pdf,'bestmodel_Rhus_NE.pdf')
dev.off()
#Now, evaluate considering bias sampling
env.files.NEwb <- list.files(path = "C:/Users/Dell/Documents/layers.NEwb", full.names = TRUE)
env.NEwb <- stack(env.files.NEwb)
env.NEwb <- setMinMax(env.NEwb)
crs(env.NEwb) <- " +proj=longlat +datum=WGS84 +ellps=WGS84 +towgs84=0,0,0"
plot(env.NEwb)
#background buffer for sampling bias
points <- read.csv("Sampling_locals.csv")[,1:2]
Sampling <- enmtools.species()
Sampling$species.name <- "Sampling"
Sampling$presence.points <- read.csv("Sampling_locals.csv")[,1:2]
Sampling$range <- background.raster.buffer(Sampling$presence.points, 25000, mask = env.NEwb)
Sampling$background.points <- background.points.buffer(points = Sampling$presence.points, radius =
25000, n = 500, mask = env.NEwb[[1]])
plot (Sampling)
plot (Sampling$background.points)
bg<-rbind (Sampling$background.points,points)
enmeval_Rhus_NEwb <- ENMevaluate(Rhus_NE_occ, env.NEwb, bg.coords = bg, method="jackknife",
RMvalues = seq(0.5, 3, 0.5), fc = c("L", "LQ"), algorithm='maxent.jar')
#Saving results
par(mfrow=c(2,2))
eval.plot(enmeval_Rhus_NEwb@results, "avg.test.AUC", variance="var.test.AUC")
dev.copy(pdf,'avgtestAUC_enmeval_Rhus_NEwb.pdf')
dev.off()
eval.plot(enmeval_Rhus_NEwb@results, "AICc")
dev.copy(pdf,'AICc_enmeval_Rhus_NEwb.pdf')
dev.off()
aic.mod <- enmeval_Rhus_NEwb@models[[which(enmeval_Rhus_NEwb@results$delta.AICc==0)]]
enmeval_Rhus_NEwb_varimpbest<-var.importance(aic.mod)
write.table(enmeval_Rhus_NEwb_varimpbest, file = "enmeval_Rhus_NEwb_varimpbest.csv")
enmeval_Rhus_NEwb_varimp<-lapply(enmeval_Rhus_NEwb@models, var.importance)
write.table(enmeval_Rhus_NEwb_varimp, file = "enmeval_Rhus_NEwb_varimp.csv")
plot(enmeval_Rhus_NEwb@predictions[[which (enmeval_Rhus_NEwb@results$delta.AICc == 0) ]])
dev.copy(pdf,'bestmodel_Rhus_NEwb.pdf')
dev.off()
```

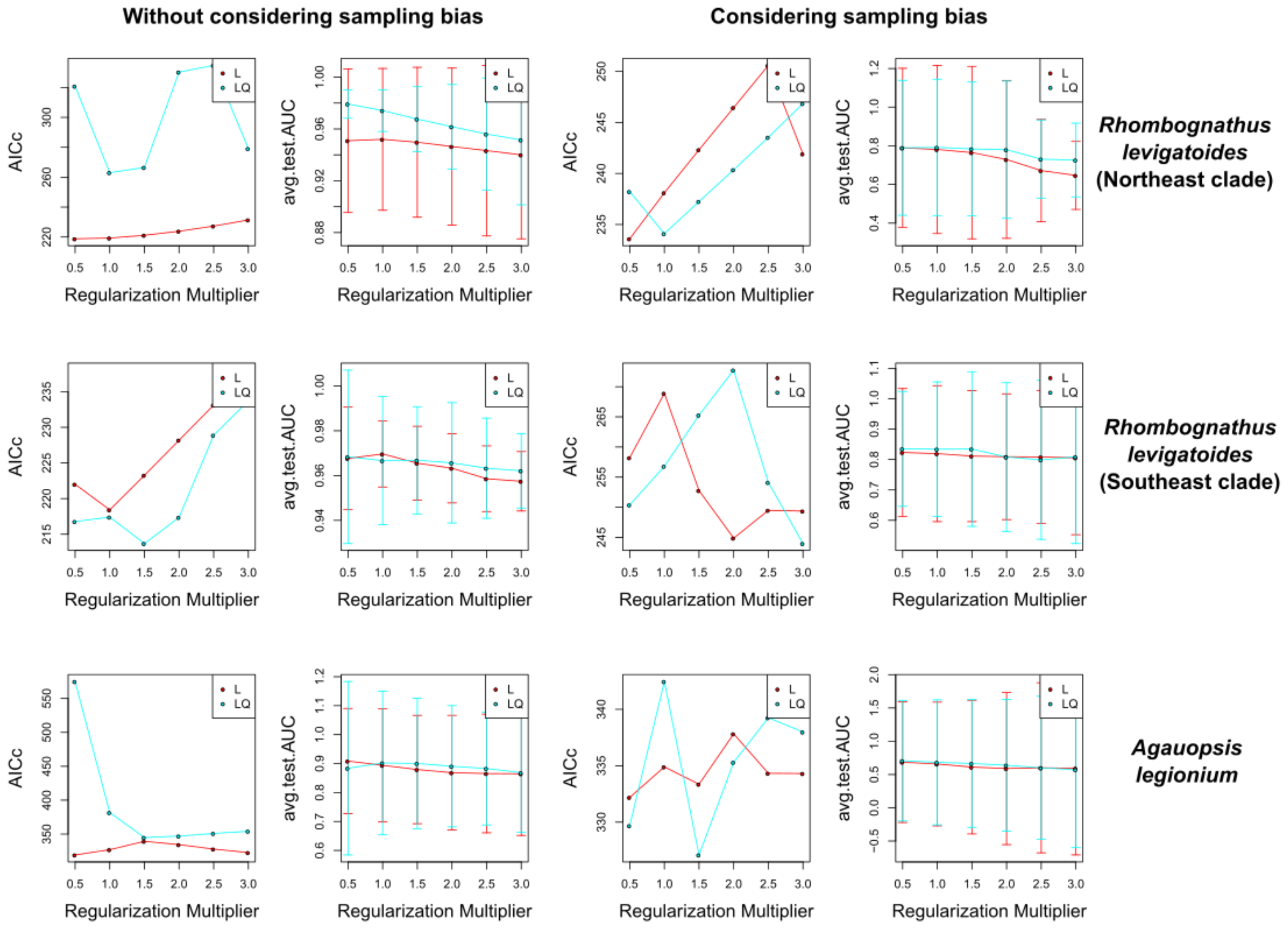


S Figure 1. ENMEval results. Plots refer to AICc values and the average of AUC Test under different values of the Regularization multiplier and Linear (red lines) and Linear and Quadratic (light blue) features.

Script 4. R script for producing a sampling bias grid for Maxent (largely taken from https://scottrinnan.wordpress.com/2015/08/31/how-to-construct-a-bias-file-with-r-for-use-in-maxent-modeling/)

```
#load libraries
library(raster)
library(MASS)
library(magrittr)
library(maptools)
#load sampling localities. Obviously change the path and names accordingly
locals <- read.csv("Sampling_locals.csv")
#load a layer to be used as mask. It must be cropped as your other enviromental layers
mask <- brick("bathy.tif")
#set projection etc.
crs(mask) <- " +proj=longlat +datum=WGS84 +ellps=WGS84 +towgs84=0,0,0"
#turn into a raster the occurrence points
occur.ras <- rasterize(locals, mask,1)
#Extract coordinates from the raster
presences <- which(values(occur.ras) == 1)
pres.locs <- coordinates(occur.ras)[presences, ]
#kde2d function gives us a two-dimensional kernel density estimate, based on the coordinates of the
#occurrence points. Note that here we set the value of bandwidth. Here we apply a very narrow area
#around sampling points. Adjust it according to your problem.
dens <- kde2d(pres.locs[,1], pres.locs[,2], 0.4, n = c(ncol(occur.ras), nrow(occur.ras)), lims = c(-52, -26, -
33, 2))
dens.ras <- raster(dens)
#Plot the raster to check
plot(dens.ras)
#Adjust values and crop using the mask. Note that a valid bias grid must have no zero or negative values.
bb <- extent(-52, -26, -33, 2)
dens.ras<- setExtent(dens.ras, bb)
masked_dens.ras<-mask(dens.ras, mask)
values(masked_dens.ras)[values(masked_dens.ras) <= 1.0e-45] = 1.0e-45
#Take a look to see if everything is ok
masked_dens.ras
plot (masked_dens.ras)
#save in a grid file. Maybe you will need to convert in an ascii file.
writeRaster(masked_dens.ras, filename="C:/Maxent/bias file.grd")
```

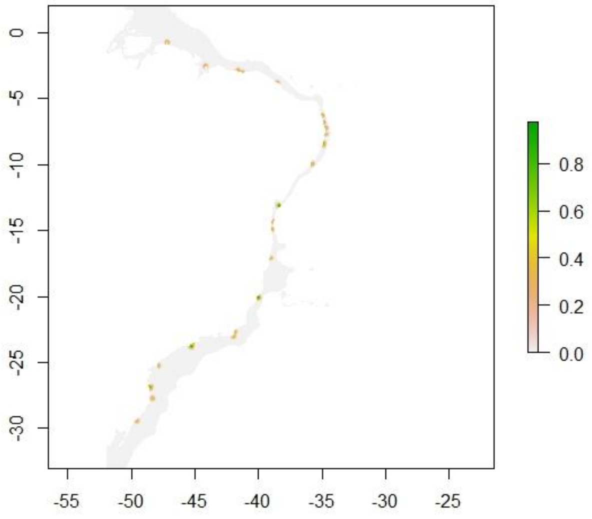


S Figure 2. Grid employed in the analyses that takes into account sampling bias.
